# Generation time is not a universal constraint on adaptive evolution

**DOI:** 10.64898/2026.02.04.703757

**Authors:** Catalina Chaparro-Pedraza, Ella Rees-Baylis

## Abstract

Conventional wisdom suggests that adaptive evolution proceeds more slowly in long-lived organisms than in short-lived ones due to longer generation times. As a result, long-lived organisms are often viewed as less capable of responding to rapid environmental change. However, empirical evidence challenges this view. Using mathematical models and demographic data from 322 wild animal populations, we show that long generation times slow adaptive evolution only under limited conditions, notably when selection acts on fecundity. When selection targets early survival, intermediate and long generation times can instead accelerate adaptive evolution. Remarkably, short-lived species tend to occupy demographic regimes in which fecundity is the dominant fitness component, whereas long-lived species occupy regimes in which early survival dominates. Therefore, both short- and long-lived species can potentially adapt rapidly, calling into question the widespread use of generation time as a general predictor of adaptive capacity to current environmental change.

## Main text

Rapid environmental change is accelerating extinction risks for biodiversity worldwide [1,2]. Understanding species’ adaptive potential is essential for enabling effective conservation efforts [3,4]. It is widely assumed that long-lived species, such as trees, large mammals, and reptiles, are especially vulnerable due to their presumed limited capacity for rapid evolutionary adaptation (e.g. ref [5–13]). This assumption is rooted in the idea that long generation times constrain the speed at which populations can respond evolutionarily to environmental change.

But does a longer generation time necessarily translate into lower adaptive potential? Despite its intuitive appeal and wide acceptance, multiple independent lines of evidence challenge this assumption. First, direct measurements of adaptive evolutionary rates do not show a decreasing trend with increasing generation time. For example, Rosenheim and Tabashnik[14] examined the relationship between the rate of insecticide resistance evolution and generation time in 682 arthropod species spanning generation times between 10 days and 2 years. They found that the rate of evolution peaks at intermediate generation times rather than in the fastest-reproducing species. A subsequent study using an expanded dataset confirmed this pattern [15]. Similarly, in a microbial evolution experiment, adaptive change has been shown to proceed as a function of absolute time rather than generation number [16].

Second, the rate of adaptation at the molecular level does not decline with generation time. In a comparative study, Rousselle et al.[17] analyzed molecular data from 50 animal species and showed that adaptive substitution rates show no such decline and may even increase slightly with longevity, which positively correlates with generation time [18,19]. A similar pattern has also been observed in long-lived plants[20]. Third, indirect evidence also suggests that the capacity for rapid adaptive evolution does not inherently differ between long- and short-lived species. When faced with environmental change, populations may adapt, go extinct, or migrate to more favorable environments. Because island species are often geographically isolated, they can only adapt or go extinct. Therefore, island ecosystems offer a unique opportunity to test whether short-lived species have a higher capacity for rapid adaptive evolution, as this should translate into a lower extinction risk compared to long-lived species. Studies of extinction risk in island birds[21,22] and reptiles[23] reveal no consistent differences between short- and long-lived species. Additionally, documented examples of fast adaptive evolution in long-lived plants [24,25], reptiles[26], birds[27], and mammals[28] are rapidly accumulating.

Together, these findings reveal a striking disconnect between a widely accepted assumption and observed evolutionary patterns: long-lived species are not inherently slow to adapt. Yet, the underlying reason for this disconnect is unclear. Here, we show that the assumption that adaptive evolution is slower in long-lived species holds only under a restricted set of conditions. Using demographic models of selection, we demonstrate that when natural selection targets survival rather than fecundity, the rate of evolution need not, and often does not, decline with increasing generation time. This insight provides a demographic mechanism explaining why long generation times do not universally constraint adaptive evolution, and why the failure to find empirical support for this constraint is not surprising but theoretically expected.

### Why doesn’t longer generation time always slow adaptive evolution?

The assumption that long-lived species are inherently slow to adapt presumes that evolution unfolds across generations. By this logic, short-lived species, which experience more generations per unit of time, are expected to adapt faster because natural selection has more frequent opportunities to act [5]. This expectation holds when selection acts on fecundity, favoring genotypes with the highest reproductive output. Since reproduction occurs more often in species with shorter generation times, the strength of selection on fecundity is expected to be stronger in these species.

However, fitness is not determined by fecundity alone. Survival often contributes more to fitness than fecundity, particularly in long-lived species [32]. Natural selection on survival operates continuously throughout the life cycle, favoring individuals that experience lower mortality at any stage. Crucially, this form of selection does not depend on the frequency of reproduction and may therefore be less sensitive to generation time.

The contrasting ways in which natural selection operates on fecundity and survival suggest that the effect of generation time on evolutionary rate depends on which fitness component is under selection. To investigate this, we use a classic demographic model of natural selection that decomposes fitness into age-specific survival and fecundity schedules, thereby linking generation time and fitness components [29–34]. The model is based on the most widely accepted measure of Darwinian fitness [29], the intrinsic growth rate *r*, implicitly defined by the continuous version of the Euler-Lotka equation,

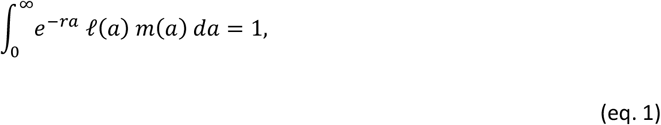

where ℓ (*a*) is the probability of surviving to age *a*, and *m(a*) is the age-specific fecundity.

Using this framework, we examine how selection acting on distinct fitness components and generation time influence the rate at which a beneficial genotype increases in frequency. According to classical population genetics, the change in frequency of a beneficial genotype A in a population otherwise composed of genotype B is

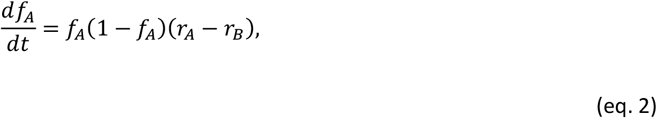

(eq. 2) where *f*_*A*_ is the frequency of the genotype A. The difference in the intrinsic growth rates of the two genotypes, (*r*_*A*_−*r*_*B*_), corresponds to the selection coefficient and determines the speed at which the beneficial genotype spreads through the population, and hence the speed of adaptive evolution.

To investigate the relationship between generation time and the speed of adaptive evolution, we use age at maturity —the average age at first reproduction— as a proxy for generation time. Although a species’ generation time depends on its survival, reproduction and maturation schedules, empirical evidence shows that only age at maturity consistently correlates with generation time across diverse plant and animal taxa [19,35]. This correlation underpins the fast–slow continuum of life history variation, with short-lived species at one end and long-lived species at the other. Moreover, in our model, reproduction and mortality rates are subject to selection, whereas age at maturity can be varied independently of selection. For these reasons, age at maturity provides a tractable and biologically interpretable axis for comparing evolutionary rates along the fast–slow continuum.

Our analytical results reveal that the relationship between age at maturity and the speed of evolution, defined by the selection coefficient, depends on the fitness component under selection (see SI1). When selection operates through fecundity, earlier maturation, and thus shorter generation times, accelerate adaptive evolution, consistent with the conventional expectation (figure 1A). In contrast, when selection targets survival, the relationship between age at maturity and evolutionary speed becomes non-monotonic and can even reverse, depending on how mortality is distributed across the life cycle.

**Figure 1.**
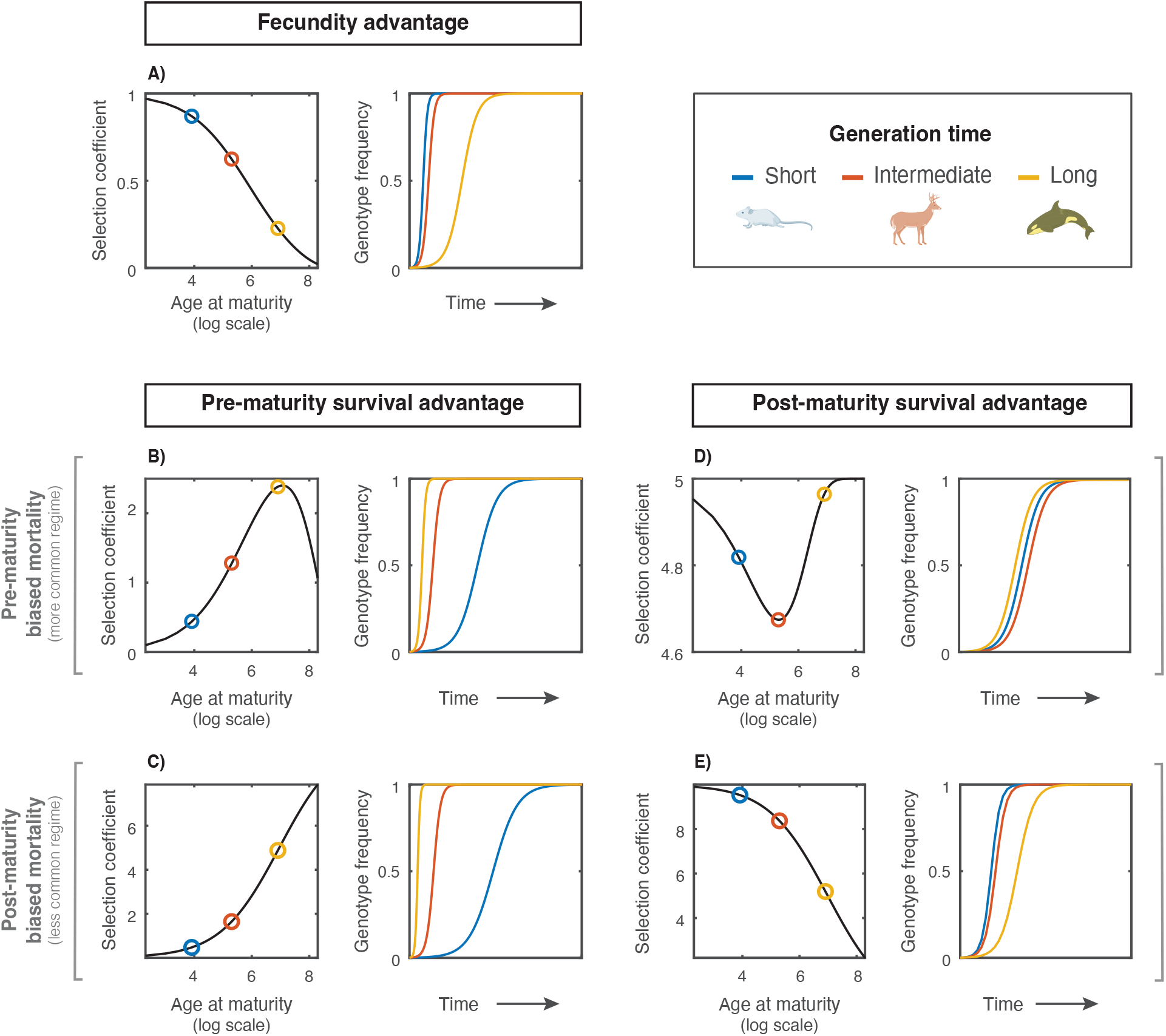
Generation time modulates the strength of selection depending on the fitness component under selection. Left panels show the selection coefficient (*r*_*A*_ *-r*_*B*_) of a rare genotype with a fitness advantage in fecundity (**A**), pre-maturity survival (**B, C**), or post-maturity survival (**D, E**) as a function of age at maturity. Right panels show the corresponding genotype frequency (*fA*) for early (blue), intermediate (red), and late (yellow) maturation. When selection acts on fecundity, the selection coefficient declines monotonically with age at maturity (**A**). When selection acts on survival, the relationship depends on the demographic regime. Under pre-maturity–biased mortality (**B, D**), selection is hump-shaped when acting on pre-maturity survival and U-shaped when acting on post-maturity survival. Under post-maturity–biased mortality (**C, E**), selection increases with age at maturity when acting on pre-maturity survival and decreases when acting on post-maturity survival. The qualitative patterns are general, and thus, independent of parameter values (see analytical investigation in SI1). For illustration, the genotype with a fitness advantage exhibits a 10% higher fecundity **(A)**, or a 10% lower mortality in the relevant stage **(B-E)**. Example early-, intermediate-, and late-maturing species are *Mus musculus* (5-8 weeks), *Odocoileus virginianus* (2 years), and *Orcinus orca* (8-12 years), respectively. An interactive version of this figure is available online (https://pcchaparro.github.io/assets/interactive_figs/InteractiveFig1.html).

In the common case where juvenile mortality exceeds adult mortality [36], selection on pre-maturity survival produces a peak in selection coefficient at intermediate ages at maturity (figure 1B). This arises because heritable differences in early survival create a filtering effect for selection: individuals with traits that enhance survival are more likely to reach reproductive maturity, while those with lower survival face an increased risk of being removed from the gene pool before reproducing. As age at maturity increases, this filter acts over a longer period, initially amplifying survival differences and strengthening selection. However, at very late maturity, further delays sharply reduce the probability of ever reaching reproduction, eventually overwhelming the survival advantage and weakening selection, thereby producing a maximum in adaptive speed at intermediate generation times. Conversely, when selection acts on post-maturity survival, the pattern is reversed: the selection coefficient initially decreases and then increases with delayed maturity (figure 1C). This is because at low ages at maturity, delaying reproduction reduces the probability of surviving to adulthood, thereby weakening the benefit of improved adult survival. But when maturity occurs late, intrinsic growth depends almost entirely on the survival of the few adults that reach this late-occurring life stage, amplifying the fitness effect of enhanced adult survival.

In the less common case where adult mortality exceeds juvenile mortality, selection on pre-maturity survival causes the selection coefficient to increase monotonically with age at maturity, because a short adult life expectancy does not dilute early survival advantages, preventing the filtering effect from weaken when age at maturity is high (figure 1D). Conversely, when selection acts on post-maturity survival, the selection coefficient declines with age at maturity because delaying maturity reduces the probability that individuals reach the mature stage, in which the adaptive benefit is realized, leading to progressively weaker selection and slower evolution (figure 1E).

These results show that the conventional expectation that long-lived species evolve more slowly holds only under limited conditions, specifically when selection acts on fecundity or when it acts on post-maturity survival in populations where mortality is concentrated after maturity. In all other cases, short-lived species do not necessarily adapt faster than other species along the fast–slow continuum. Whether this occurs depends on the life stage in which the fitness advantage arises and on the distribution of mortality across the life cycle.

These findings raise the question: how do these different fitness components contribute to the intrinsic growth rate of natural populations, and how does this contribution vary with age at maturity? To address this, we leverage standardized demographic matrices derived from field-based life-history data in the COMADRE Animal Matrix Database [37] to assemble a comparative dataset from 322 wild populations representing 129 species spanning diverse animal taxa (see Methods and figure S1). For each population, we quantified the relative contributions of fecundity, pre-maturity and post-maturity survival to growth rate using elasticity analysis, a widely used tool for identifying the demographic rates to which fitness is most sensitive [38].

Across species, we find contrasting patterns among fitness components (figure 2). The contribution of fecundity declines with increasing age at maturity, whereas the contribution of pre-maturity survival increases, becoming more important in species that delay reproduction. In contrast, the influence of post-maturity survival remains roughly constant across the observed range of maturation ages, indicating that changes in adult survival affect fitness to a similar degree in both short- and long-lived species. A similar pattern was previously reported for mammals alone [39]; our broader analysis shows that it generalizes across animal taxa. These results suggest that even small improvements in pre-maturity survival can generate large increases in intrinsic growth rate for late-maturing species, potentially enabling rapid adaptive responses when selection targets early survival. Thus, along the fast–slow continuum, the fitness component contributing most to intrinsic growth rate shifts systematically— from fecundity in fast-living species to pre-maturity survival in the slow-living species.

**Figure 2.**
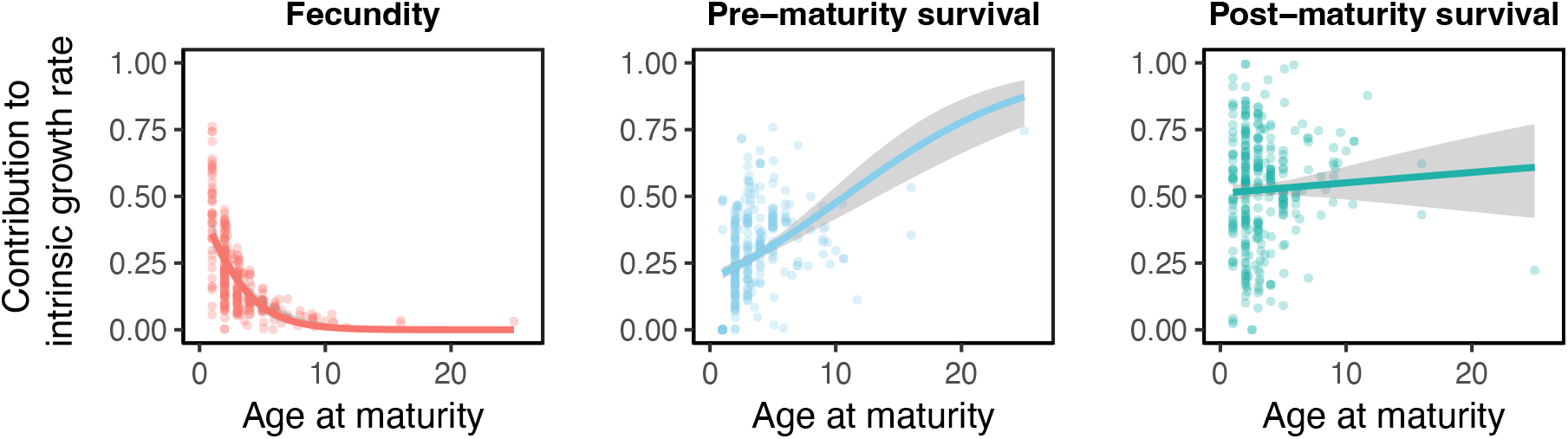
Contributions of different fitness components to growth rate in wild populations. The relative contribution of fecundity **(A)**, pre-maturity survival **(B)** and post-maturity survival **(C)** to the intrinsic growth rate as a function of the age at maturity for 322 wild populations from 129 animal species. Contributions are quantified using elasticity analysis applied to each population’s projection matrix. Lines show fitted trends (using generalised linear models with quasibinomial errors and logit link), with shaded bands indicating the 95% confidence intervals.

The empirical patterns shed light into how our analytical results may play out in nature. In fast-living species, the dominance of fecundity as a fitness component places populations in a demographic regime in which our model shows that when selection targets this fitness component adaptive change can be the fastest. Conversely, in slow-living species, the dominance of pre-maturity survival corresponds to the regime in which selection on early survival can yield the fastest adaptive responses. Along the fast–slow continuum, the fitness component contributing most to intrinsic growth rate therefore coincides with the fitness component through which selection can act most efficiently, allowing high potential rates of adaptive evolution along the continuum.

### Adaptive potential under environmental change

If long generation times do not necessarily slow adaptive evolution, what does this imply for biodiversity facing rapid environmental change? This question is particularly pressing in the current context of climate change, where long-lived species are often assumed to have inherently limited adaptive capacity in climate vulnerability assessments [40–42]. Global temperatures have been rising at a rate of 0.2°C per decade since the 1970s [43], imposing sustained directional selection across taxa with widely varying life histories.

Crucially, this selection pressure unfolds in absolute time. As a result, long- and short-lived species experience the same environmental change over very different numbers of generations. Understanding how species with contrasting generation times respond to such gradual but persistent change is therefore essential. Because contemporary adaptive evolution typically involves many loci of small effect rather than single mutations [20], we adopt a quantitative genetic framework. At the same time, environmental stress can drive populations to low sizes, making demographic stochasticity unavoidable. Under these conditions, genetic drift and extinction risk become integral components of the evolutionary dynamics, interacting directly with selection.

To capture these complex dynamics, we simulated adaptation in populations facing a gradually changing environment, such as rising temperatures, using an individual-based model. Environmental change imposes sustained directional selection on a quantitative trait influencing different fitness components in separate scenarios: fecundity, pre-maturity survival, and post-maturity survival (figure 3 A-C). Individuals whose trait value more closely match the shifting optimum experience higher birth rate or lower mortality rate, depending on the scenario. Population size emerges dynamically from the underlying birth and death schedules. Because contemporary adaptation in natural populations often relies on standing genetic variation [42], we focus on evolutionary responses driven by selection and drift, excluding mutation.

**Figure 3.**
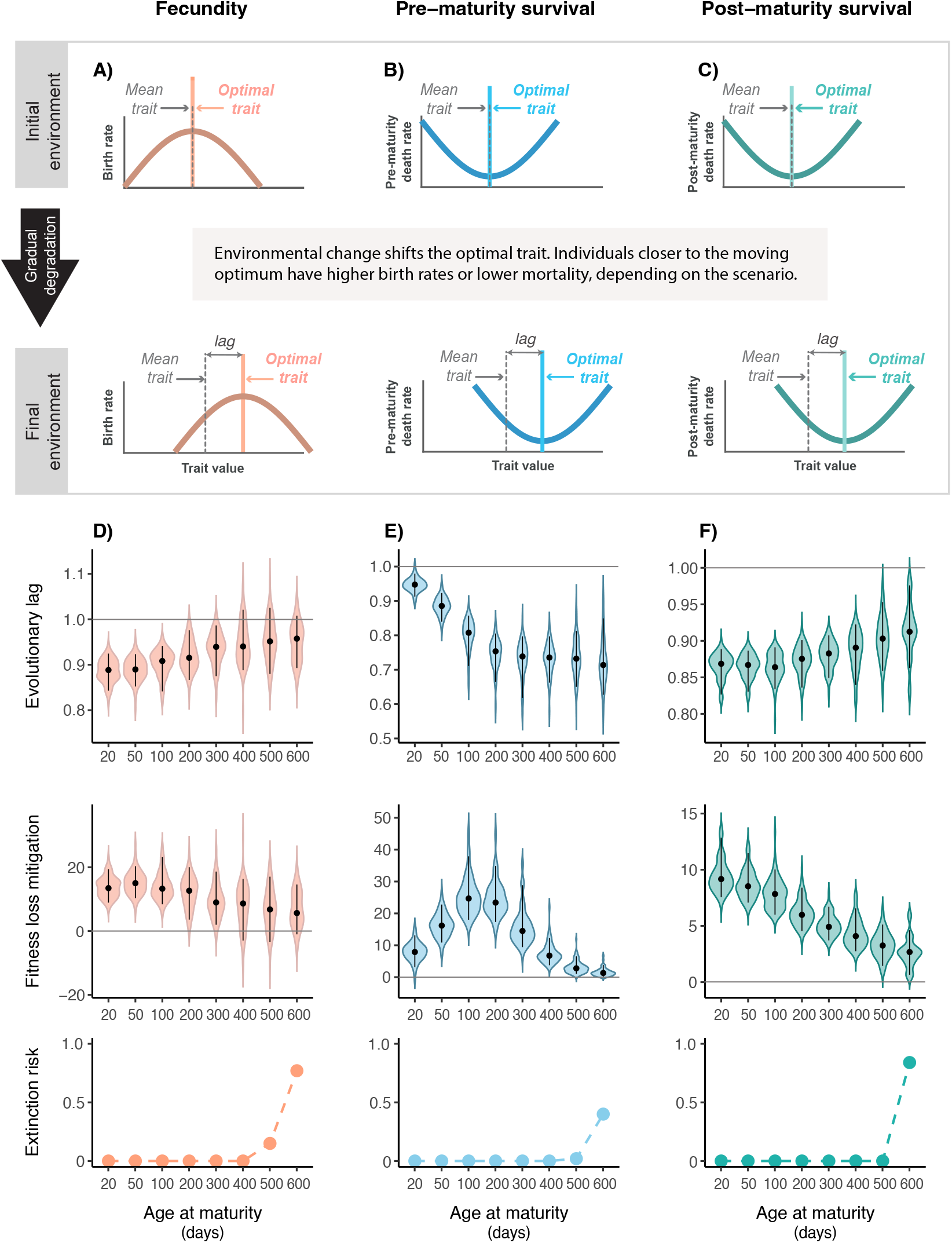
Adaptive responses to gradual environmental change with fixed demographic rates. The individual-based model structure is illustrated in the top panels (grey box). Individuals closer to the shifting optimum experience higher fecundity (**A**), lower pre-maturity mortality (**B**), or lower post-maturity mortality (**C**). Bottom panels show evolutionary lag (**D–F**, top row), fitness loss mitigated by evolution (middle row), and extinction risk (bottom row) as functions of age at maturity. Populations differ only in age at maturity. Each violin represents the distribution of 100 replicate simulations; black dots indicate means, and black lines SD. Evolutionary lag values of 1 (horizontal grey line) indicate no change relative to the initial phenotype, while values below or above 1 indicate adaptive tracking or maladaptive evolutionary responses, respectively. Fitness-loss mitigation is expressed as the percentage reduction in fitness loss relative to a non-evolving population. Negative values of fitness loss mitigation arise when evolutionary responses were maladaptive. The extinction risk corresponds to the fraction of simulations in which the population went extinct. Parameter values as in table SI2.2.

Adaptive capacity in each simulation is characterized using two complementary measures: evolutionary lag quantifies how closely the population mean trait tracks the shifting optimum, whereas fitness loss mitigation captures the demographic benefit of adaptation in terms of preserved intrinsic growth rate. Under a simplified demographic framework, in which only age at maturity varies, simulations show that the relationship between adaptive capacity and age at maturity depends on which fitness component is under selection (figure 3 D-F). When selection acts through fecundity or post-maturity survival, the evolutionary lag increases steadily with age at maturity, indicating that short-lived species adapt faster, and this enables a higher mitigation of fitness loss. In contrast, when selection operates through pre-maturity survival, early-maturing species exhibit the largest evolutionary lag. This occurs because a longer pre-maturity period allows selection to remove individuals with higher intrinsic juvenile mortality more efficiently, consistent with our analytical results. However, this increased efficiency does not directly translate into higher mitigation of fitness loss: in very late-maturing species, improvements in daily survival become diluted across the extended pre-reproductive period and, thus, contribute little to overall fitness. As a result, fitness-loss mitigation peaks at intermediate rather than late maturity. Across all scenarios, late-maturing species have smaller population sizes, which strengthens genetic drift and increases their vulnerability to extinction.

Because wild populations exhibit strong covariation between demographic rates and age at maturity, we next repeated the simulations using empirically parameterized life-history schedules (figure 4). These simulations confirmed the qualitative patterns from the simplified demographic framework for the evolutionary lag and the fitness-loss mitigation when selection targeted fecundity and post-maturity survival. However, when selection acted on pre-maturity survival, the evolutionary lag followed a U-shaped relationship with age at maturity, reaching its minimum at intermediate age at maturity and, thus, mirroring the non-monotonic pattern predicted by the analytical model. In contrast to the simplified demographic case, extinction risk showed no consistent relationship with age at maturity because the covariation of late maturation with other demographic rates buffers population size. Indeed, late-maturing species consistently exhibit very low mortality (e.g. the Yellow Mud Turtle, *Kinosternon flavescens*), or high fecundity (e.g. piscivorous fish like the Northern Pike, *Esox lucius*), reducing their susceptibility to small population sizes, and consequently genetic drift and extinction.

**Figure 4.**
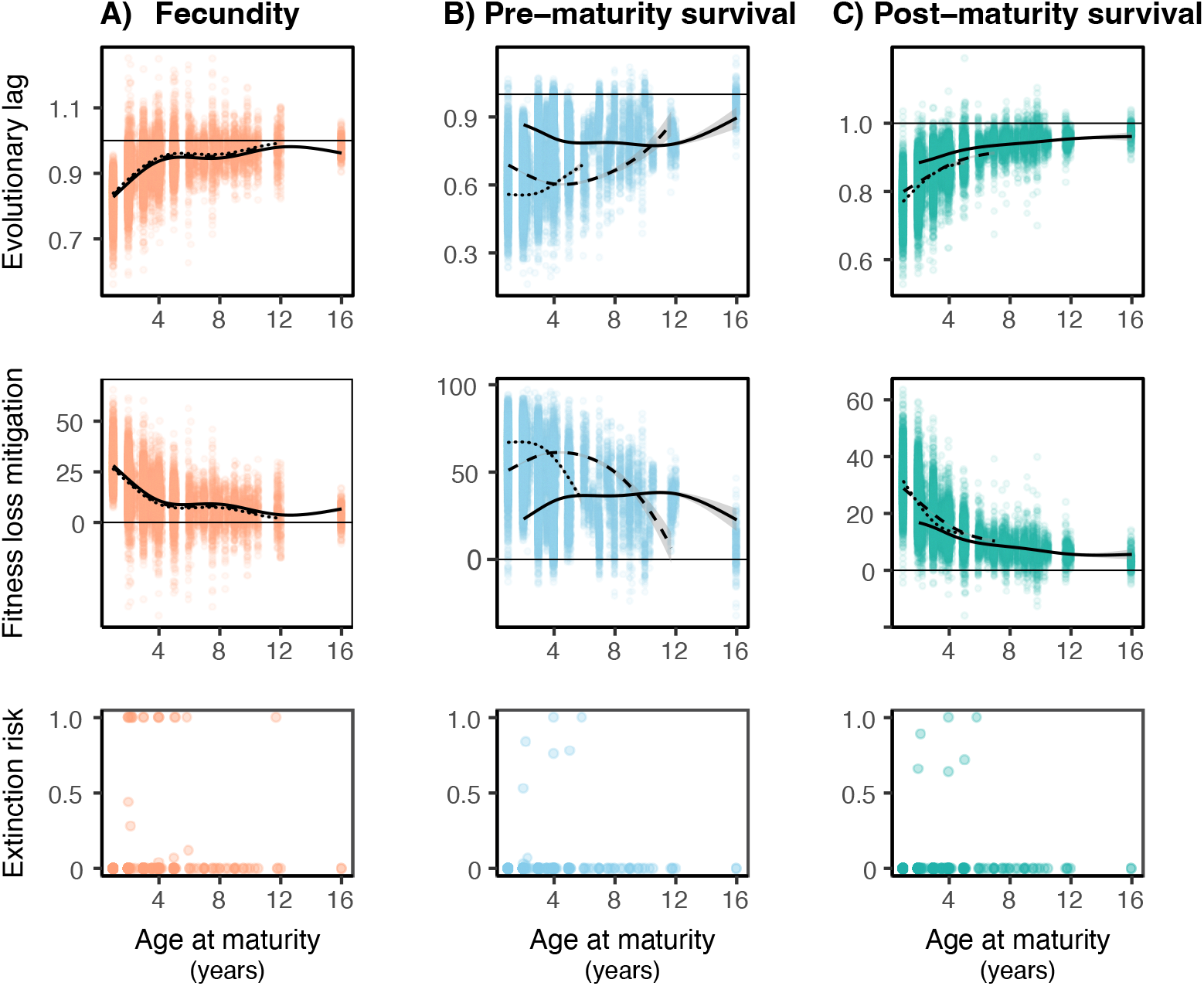
Adaptive responses to gradual environmental change using empirical demographic data. Evolutionary lag (top row), fitness loss mitigated by evolution (middle row), and extinction risk (bottom row) are shown as functions of age at maturity when selection acts on fecundity **(A)**, pre-maturity survival **(B)**, or post-maturity survival **(C)**. Each point represents the outcome of one of 100 simulations for each of 277 empirically parameterized populations. Black lines show GAM fits and shaded ribbons indicate 95% confidence intervals. Because baseline fecundity or mortality covaries with age at maturity (see figure S2), populations were grouped into low (solid line), intermediate (dashed line), and high (dotted line) baseline classes using the 30th and 70th percentiles. Parameter values as in table SI2.3.

Together, these two analyses provide complementary insight into how generation time influences the ability of populations to adapt to a gradually changing environment. While the simplified demographic analysis isolates the causal effects of age at maturity and the fitness component under selection, the empirically parameterized analysis allows assessing whether the theoretical predicted patterns are robust once realistic demographic covariation is introduced. Their combined results show that longer generation times neither necessarily elevate extinction risk nor universally slow adaptation, revealing that generation time alone is a poor predictor of adaptive capacity and extinction risk.

## Discussion

Our results demonstrate that generation time is not a universal constraint on adaptive evolution, contrary to the widespread assumption that long-lived species are inherently slow to adapt. Many studies that invoke this assumption actually refer to molecular evolution rates, such as DNA substitution rates, rather than rates of adaptive evolution. The two are often conflated, but they are not equivalent. Molecular evolution studies typically measure neutral substitution rates (e.g., synonymous mutations). These rates are influenced by generation time because short-lived species undergo more DNA replications per unit time, leading to higher mutation rates, as it has been documented in plants[44] and animals [45–50]. However, higher mutation rates do not necessarily translate into faster adaptation, which also depends on selection pressures and genetic architecture [51]. For instance, in long-term experimental evolution, *E. coli* populations evolving over 60,000 generations showed decelerated adaptation despite constant molecular evolution [52]. Therefore, molecular evolution rates cannot be used as a general proxy for adaptive speed.

We show that the conventional expectation that shorter generation times accelerate adaptation holds when selection acts on fecundity, where reproductive turnover sets the pace of evolution. When selection instead targets pre-maturity survival, our model predicts a fundamentally different outcome: the speed of adaptation initially increases and then declines with age at maturity (figure 1B). Strikingly, the evolution of pesticide resistance in arthropod pests exhibits the same pattern [14,15], which until now lacked a mechanistic explanation [15]. Our results provide such an explanation: when selection acts on pre-maturity survival, adaptation proceeds by increasing the probability of reaching reproductive age, creating a filtering effect that strengthens as the pre-reproductive period lengthens. However, at very late maturity, further reproductive delay sharply reduces the probability of reproducing at all, eventually overwhelming the survival advantage and weakening selection, thereby causing adaptive speed to peak at intermediate generation times.

These results indicate that the relationship between generation time and adaptive evolution rate depends critically on how selection is distributed across fitness components. Our comparative demographic analysis shows that the relative importance of these components shifts systematically along the fast–slow continuum, with the dominant contribution to fitness shifting from fecundity in fast-living species to pre-maturity survival in slow-living species. This redistribution along the fast–slow continuum is consistent with genomic evidence showing that evolutionary transitions to fast life histories are accompanied by strengthened selection on reproductive functions and relaxed selection on aging- and maintenance-related genes [53].

Our findings underscore the role of survival-driven selection, especially during pre-reproductive life stages, in shaping evolutionary trajectories of long-lived species. Yet such selection has often been overlooked because many theoretical models fail to incorporate realistic organismal life history. Capturing the filtering effect described above requires explicitly considering immature stages. However, most existing models in evolutionary ecology neglect this aspect (reviewed in [54], but see [55,56]), implicitly assuming that individuals mature instantly after birth and, thus, obscuring the role of pre-maturity survival in evolutionary dynamics. This limitation is compounded in foundational evolutionary frameworks, such as the Wright–Fisher model, which assume non-overlapping generations and thus collapse the entire life cycle into a single reproductive event per generation. Under these assumptions, survival becomes mathematically indistinguishable from fecundity, and all selection is forced to act through per-generation reproductive output. These structural simplifications persist in many more recent models, even when generation time is incorporated explicitly (e.g. [57–59]).

Our models consider scenarios in which selection acts predominantly through a single fitness component. This simplification allows us to isolate how selection operating through different fitness components interacts with life history to shape evolutionary dynamics. In natural populations, however, quantitative traits often influence multiple fitness components simultaneously. In such cases, evolutionary responses are expected to reflect weighted combinations of the patterns described here, although future work will be needed to determine how interactions among fitness components modify these dynamics. Likewise, our simulations consider comparable levels of standing genetic variation along the fast–slow life-history continuum. Although standing genetic variation may in principle covary with life history through differences in effective population size, available empirical estimates do not reveal a consistent relationship between genetic variation in fitness and position along the fast–slow continuum [60]. As a result, standing genetic variation is not expected to systematically favor either fast- or slow-living species and therefore does not affect our qualitative findings.

By linking life-history structure to evolutionary dynamics, our study identifies a demographic mechanism whereby selection acting on survival can drive rapid adaptation, even in long-lived species. This mechanism explains previously unresolved empirical patterns and shows that generation time alone is a poor predictor of adaptive capacity. Our results therefore caution against using generation time as a proxy for adaptive potential in assessments of species’ vulnerability to rapid environmental change and underscore the need for demographically informed models to generate accurate predictions that enable more effective conservation strategies.

## Methods

### Quantifying the contribution of different fitness components to the intrinsic growth rate in natural populations

Empirical demographic data were obtained from the COMADRE database version 4.23.3.1 [37] and analysed in R version 4.4.0 using the packages *Rcompadre* and *Rage[61]*. We included only pooled or mean matrices that met the following criteria: no missing values; ergodic, irreducible, and primitive; unmanipulated treatments; projection interval of 1 year; decomposable into survival and fecundity matrices; matrix dimension ≥ 2×2; non-zero fecundity; and derived from wild populations. After filtering, 333 matrices from 136 species remained. Matrices were then manually screened to ensure biologically interpretable estimates of age at maturity, resulting in exclusion of 11 observations and adjustment of the default life starting stage for 10 cases; specific justifications are documented in table S1. The final dataset comprised 322 matrices from 129 species, but results were qualitatively unchanged when analyses were repeated using the complete dataset (333 matrices) and default starting stage.

For each matrix, age at maturity was estimated as the mean age at first reproduction using the *mature_age* function in *Rage*. Elasticity analyses were conducted with *perturb_matrix* from *Rage*. Fecundity elasticities included contributions from both the sexual and clonal reproduction (although only one species exhibited clonal reproduction, the Red tree sponge, *Amphimendon compressa*). To distinguish between pre-maturity and post-maturity survival, we classified reproductive stages as those with non-zero column sums in the fecundity matrix, and all earlier stages as pre-reproductive. Survival elasticities were then summed within each life history stage to obtain the contribution of pre-maturity and post-maturity survival to the intrinsic growth rate.

### Adaptation in populations facing a gradually changing environment using Individual based-model simulations

We implement an individual-based model that explicitly tracks survival, reproduction, and trait inheritance through time. The simulation proceeds in discrete time steps Δ*t*, interpreted as units of absolute time (days), rather than generations. We simulate three distinct scenarios, each defined by the fitness component through which a quantitative trait influences fitness: fecundity, pre-maturity survival, or post-maturity survival.

To model a gradually deteriorating environment, we allow the optimal phenotype *θ* to shift at a constant rate ϵ, such that Δ*θ*=ϵ Δ*t*. In each scenario, an individual ‘s fitness component under selection depends on the mismatch between its trait *z* and the optimal phenotype *θ*. When selection acts on fecundity, the mismatch reduces reproductive output, whereas when selection acts on survival, the mismatch increases either juvenile or adult mortality (figure 3). Model details are provided in SI2.

We simulate 50 years of gradual environmental change, during which *θ* shifts continuously, followed by 20 years in which the optimum remains constant. This second phase is required to obtain meaningful estimates of extinction risk. For each species or parameter set, we simulate 100 replicate trajectories. From each replicate, we compute two measures of adaptive performance: the evolutionary lag and the loss of fitness mitigated by evolution. The evolutionary lag is defined as the difference between the population mean trait and the final optimum, scaled by the optimum, such that lag = 0 indicates perfect tracking and lag = 1 indicates no evolutionary change. Fitness loss mitigation quantifies the extent to which evolution compensates for the fitness decline caused by environmental change and is calculated as:

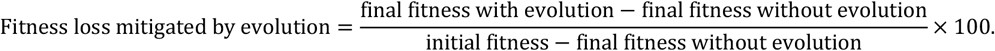

Here, initial fitness is the mean fitness at the start of the simulation (when the population is optimally adapted), final fitness without evolution is the mean fitness that the initial population would experience in the fully degraded environment (i.e., after *θ* reaches its maximum), and final fitness with evolution is the mean fitness of the evolved population at the end of the simulation, after *θ* has reached the same maximum value.

Fitness is quantified using the intrinsic growth rate as defined by the continuous version of the Euler-Lotka equation.

## Acknowledgements

The authors thank Andreas Scheidegger for making the interactive version of Figure 1, as well as Philine Feulner, Lotte de Vries and Gregory Roth for helpful feedback on an early version of this manuscript.

## Funding

CCP was supported by the SNF Ambizione Grant No 223497.

## Author contributions

Conceptualization: C.C.P., Methodology: C.C.P., E.R.B, Analysis: C.C.P., E.R.B, Visualization: C.C.P., E.R.B. Writing—original draft: C.C.P. Writing—review & editing: C.C.P., E.R.B.

## Competing interests

The authors declare that they have no competing interests.

## Data, code, and materials availability

Demographic data from wild animal populations were obtained from published studies (as cited in the text). Code for running simulations and reproducing analyses presented in this paper will be hosted at Zenodo. For revision purposes, we provide the code as a supplementary material.

---

**Figure S1.**
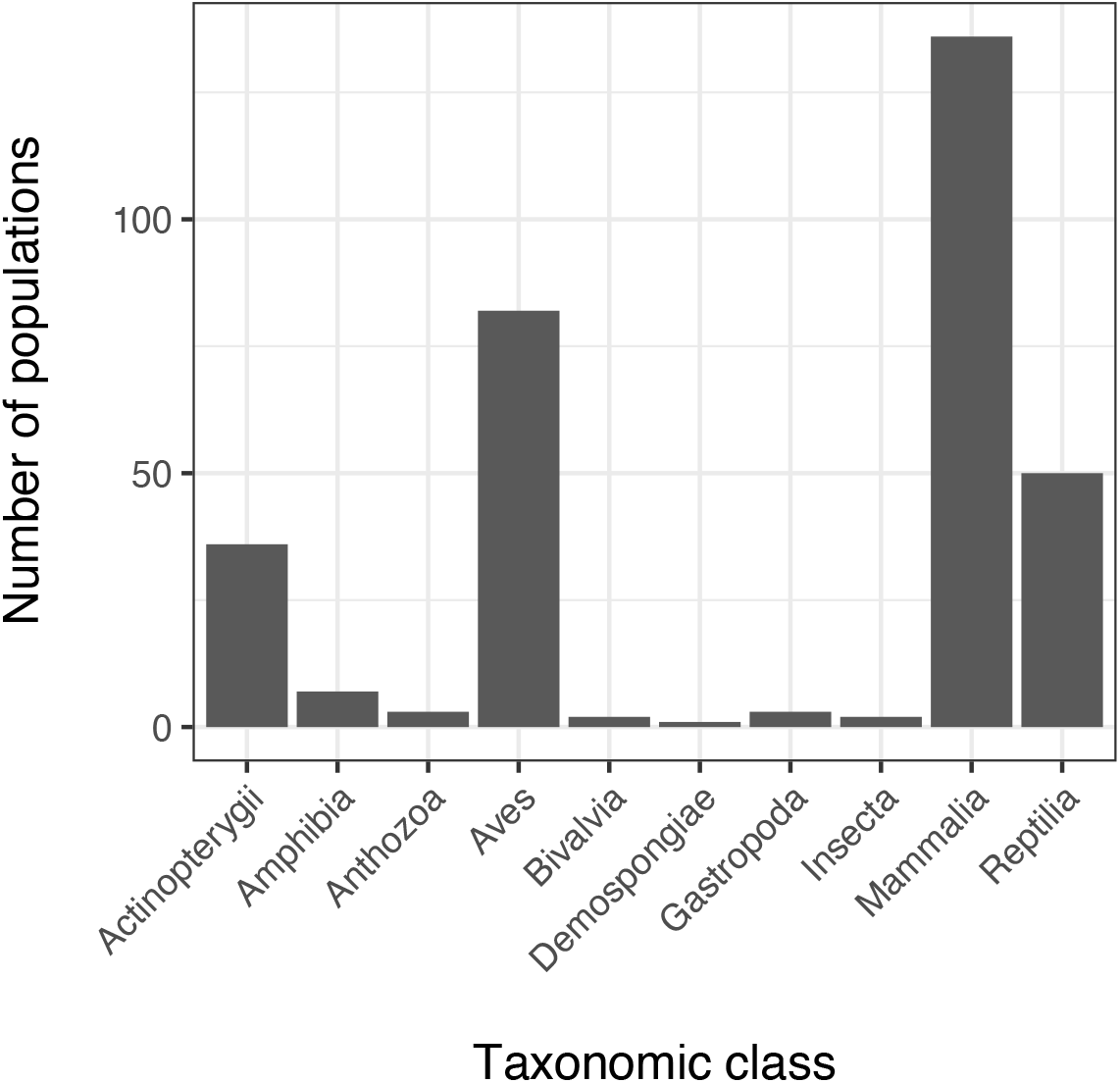
Taxonomic distribution of populations included in the comparative demographic analysis. Bars show the number of populations per taxonomic class included (total n = 322 populations across 129 species). The dataset spans major vertebrate and invertebrate clades, with strongest representation of Aves and Mammalia.

**Figure S2.**
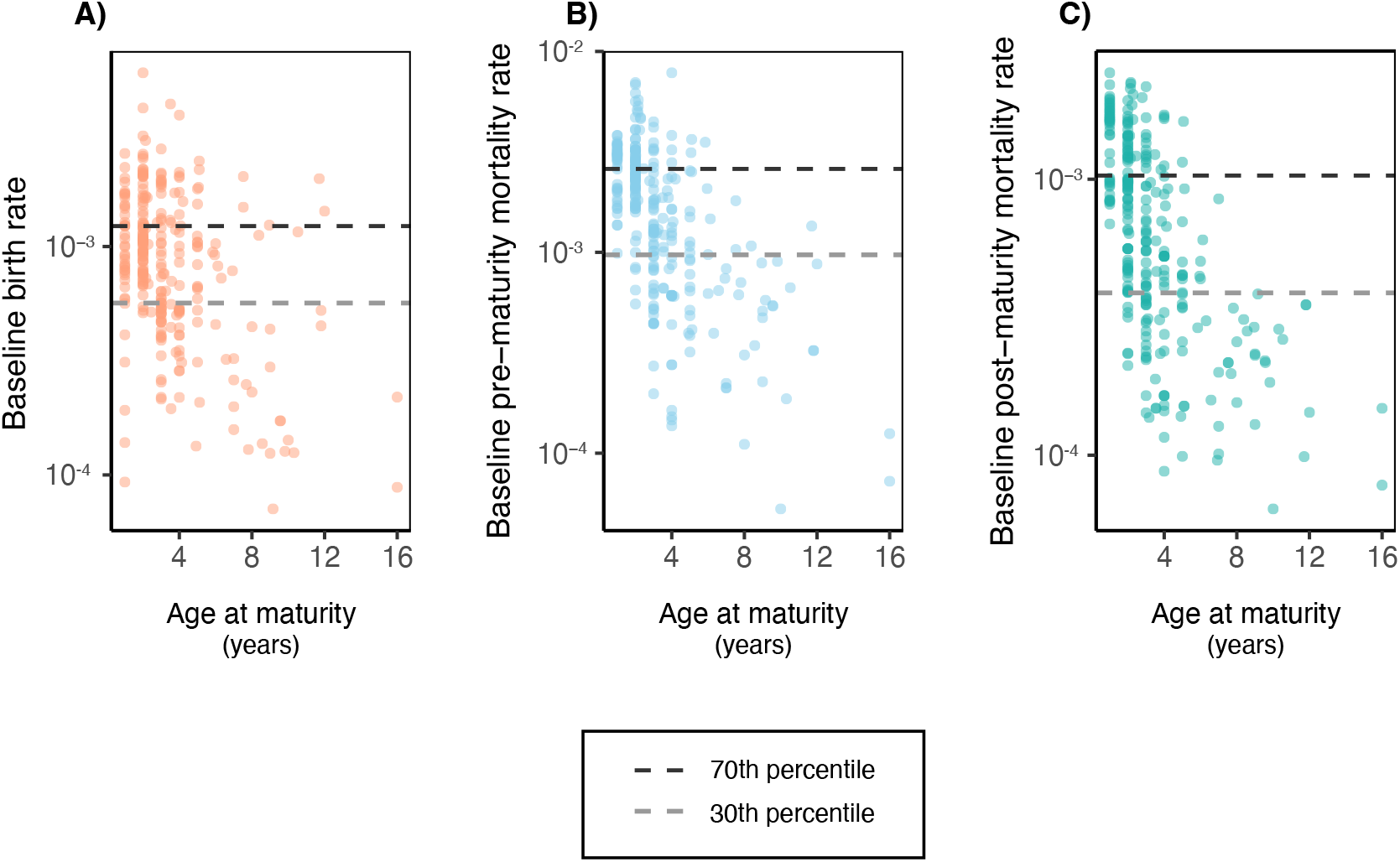
Baseline demographic rates across populations used to parameterize the Individual-based model. Empirical relationship between age at maturity and baseline vital rates used to parameterize the simulations for birth rate (**A**), pre-maturity (**B**) and post-maturity mortality rates (**C**) (n = 277 populations). Each point represents one population’s estimated baseline rate on a logarithmic scale. Horizontal dashed lines indicate the 30th and 70th percentiles of each rate distribution, which were used to define the low, medium, and high demographic-rate classes in the GAM analyses. Mortality rates exhibit a stronger covariation with age at maturity than birth rates.

## Notes

### Competing Interest Statement

The authors have declared no competing interest.

