## Supplementary material for "Generation time is not a universal constraint on adaptive evolution"

This supplementary note includes:

- Supplementary text:
    1. Analytical investigation of selection coefficient for different fitness components
    2. Description of the Individual-Based Model
- Supplementary figures:
    1. Figure S1: Taxonomic distribution of populations included in the comparative demographic analysis.
    2. Figure S2: Baseline demographic rates across populations used to parameterize the Individual-based model.
- Supplementary table:
    1. Table S1: Metadata of matrices dataset used in figure 2.

Other supplementary materials for this manuscript includes: MATLAB code of the model for IBM simulations and R code for result visualization.

### Supplementary Information 1. Analytical investigation of selection coefficient for different fitness components

Here, we investigate how selection acting on distinct fitness components (i.e. fecundity, pre-maturity mortality, and post-maturity mortality) influences the rate at which a beneficial genotype increases in frequency, and how this rate depends on age at maturity. According to classical population genetics models, the rate of change of genotype frequency is proportional to the selection coefficient (see eq. 2). We therefore examine how the selection coefficient is affected by age at maturity when selection acts on distinct fitness components. We consider two genotypes: genotype A, which has a fitness advantage, and emerges in a population dominated by genotype B. The two genotypes have fecundity and mortality schedules determined by age $a$, and characterized by their birth rate $\beta_i$, pre-maturity mortality rate $\delta_i$, and post-maturity mortality rate $\mu_i$, where $i \in \{A, B\}$. Both genotypes mature at age $a_m$. Using the Euler Lotka equation (eq. 1), we derive an expression for the intrinsic growth rate $r_i$ of genotype $i$, which we then use to calculate the selection coefficient, $c$, in three scenarios exploring the effects of distinct fitness components. All rates $(\beta_i, \mu_i, \delta_i, r_i)$ have units of time$^{-1}$ and $a_m$ units of time.

**Derivation of the intrinsic growth rate and the selection coefficient** To obtain an analytical expression for the intrinsic growth rate $r_i$ of genotype $i$, we use the Euler–Lotka equation:

$$\int_0^\infty e^{-r_i a} l_i(a) m_i(a)\, da = 1, \tag{SI1.1}$$

where

$$l_i(a) = \begin{cases} e^{-\delta_i a}, & a < a_m, \\ e^{-\delta_i a_m} e^{-\mu_i (a - a_m)}, & a \geq a_m, \end{cases} \qquad m_i(a) = \begin{cases} 0, & a < a_m, \\ \beta_i, & a \geq a_m. \end{cases}$$

Substituting $l_i(a)$ and $m_i(a)$, yields

$$\beta_i e^{-\delta_i a_m} e^{\mu_i a_m} \int_{a_m}^\infty e^{-(r_i + \mu_i) a}\, da = 1,$$

which, after evaluating the integral, can be rewritten as

$$\beta_i \frac{e^{-(r_i + \delta_i) a_m}}{r_i + \mu_i} = 1.$$

Then, we rearrange for $r_i$ to obtain the implicit equation

$$\beta_i = (r_i + \mu_i)\, e^{(r_i + \delta_i) a_m}. \tag{SI1.2}$$

We define $x_i = r_i + \mu_i$ and $\Delta_i = \delta_i - \mu_i$ and rewrite (SI1.2) as

$$\begin{aligned} \beta_i &= x_i e^{(x_i + \Delta_i) a_m} \\ \beta_i e^{-a_m \Delta_i} &= x_i e^{x_i a_m} \end{aligned} \tag{SI1.3}$$

Note that when juvenile mortality exceeds adult mortality ($\delta_i > \mu_i$), $\Delta_i > 0$; conversely, when adults have a higher mortality than juveniles, $\Delta_i < 0$.

Multiplying both sides by $a_m$ and applying the definition of the Lambert $W$ function ($ye^y = z \Rightarrow y = W(z)$) yields

$$a_m x_i = W\left(a_m \beta_i e^{-\Delta_i a_m}\right). \tag{SI1.4}$$

When substituting $x_i$ and $\Delta_i$ in this equation, it follows that

$$a_m(r_i + \mu_i) = W\left(a_m \beta_i e^{-a_m(\delta_i - \mu_i)}\right), \tag{SI1.5}$$

which can be solved for $r_i$, yielding

$$r_i = \frac{1}{a_m} W\left(a_m \beta_i e^{a_m(\mu_i - \delta_i)}\right) - \mu_i. \tag{SI1.6}$$

The selection coefficient, defined as the difference in growth rates, is thus given by

$$c = r_A - r_B = \frac{1}{a_m}\left[\left(W\left(a_m \beta_A e^{a_m(\mu_A - \delta_A)}\right) - \mu_A\right) - \left(W\left(a_m \beta_B e^{a_m(\mu_B - \delta_B)}\right) - \mu_B\right)\right]. \tag{SI1.7}$$

### 1 Scenario. Fitness advantage in fecundity

In the first scenario, we consider the case in which genotype A has a fitness advantage in fecundity, such that $\beta_A > \beta_B$, whereas all other demographic rates are equal. Hence, $\delta_A = \delta_B = \delta$ and $\mu_A = \mu_B = \mu$. In such a case, the selection coefficient is

$$c_f = \left(\frac{W_A}{a_m} - \mu\right) - \left(\frac{W_B}{a_m} - \mu\right) = \frac{1}{a_m}\left(W_A - W_B\right) \tag{SI1.8}$$

where $W_A = W\left(a_m \beta_A e^{a_m(\mu-\delta)}\right)$ and $W_B = W\left(a_m \beta_B e^{a_m(\mu-\delta)}\right)$. To understand how $c_f$ depends on $a_m$, we calculate the derivative of this equation with respect to $a_m$

$$\frac{\partial c_f}{\partial a_m} = -\frac{1}{a_m^2}\left(W_A - W_B\right) + \frac{1}{a_m}\left(\frac{\partial W_A}{\partial a_m} - \frac{\partial W_B}{\partial a_m}\right). \tag{SI1.9}$$

The derivative of the Lambert $W$ function satisfies

$$\frac{dW(x)}{dx} = \frac{W(x)}{x\left[1 + W(x)\right]}. \tag{SI1.10}$$

Hence,

$$\frac{\partial W_i}{\partial a_m} = \frac{W_i}{z_i(1 + W_i)} \frac{\partial z_i}{\partial a_m}.$$

where $z_i(a_m) = a_m\, \beta_i\, e^{a_m(\mu-\delta)}$, and

$$\frac{\partial z_i}{\partial a_m} = \beta_i e^{a_m(\mu-\delta)}\left[1 + (\mu - \delta)a_m\right] = \frac{z_i}{a_m}\left[1 + (\mu - \delta)a_m\right].$$

Therefore,

$$\frac{\partial W_i}{\partial a_m} = \frac{W_i}{1 + W_i} \frac{1 + (\mu - \delta)a_m}{a_m}.$$

Substituting into the equation (SI1.9) gives

$$\begin{aligned}
\frac{\partial c_f}{\partial a_m} &= \frac{1}{a_m^2}\left[-\left(W_A - W_B\right) + \left(1 + (\mu - \delta)a_m\right)\left(\frac{W_A}{1 + W_A} - \frac{W_B}{1 + W_B}\right)\right] \\
&= \frac{1}{a_m^2}\left(W_A - W_B\right)\left[-1 + \left(1 + (\mu - \delta)a_m\right)\left(\frac{1}{(1 + W_A)(1 + W_B)}\right)\right] \\
&= \frac{1}{a_m^2}\left(W_A - W_B\right)\left[\frac{a_m(\mu - \delta) + 1}{(W_A + 1)(W_B + 1)} - 1\right]
\end{aligned} \tag{SI1.11}$$

We know that:

1. $a_m > 0$, therefore $\frac{1}{a_m^2} > 0$

2. $\beta_A > \beta_B$, therefore $W_A > W_B$

3. $W_A > W_B > 0$, therefore $W_A - W_B > 0$, and $(W_A + 1)(W_B + 1) > 1$

Therefore, the term in the squared brackets determines the sign of the equation, and thus how the selection coefficient varies with age at maturity. This term is negative when $a_m(\mu - \delta) + 1 < (W_A + 1)(W_B + 1)$. In the following, we examine under which conditions this inequality is satisfied.

$$\begin{aligned}
a_m(\mu - \delta) + 1 &< (W_A + 1)(W_B + 1) \\
a_m(\mu - \delta) + 1 &< W_A + W_B + W_A W_B + 1 \\
a_m(\mu - \delta) &< W_A + W_B + W_A W_B \\
a_m(\mu - \delta) &< a_m(r_A + \mu) + a_m(r_B + \mu) + a_m^2(r_A + \mu)(r_B + \mu) \\
\mu &< r_A + r_B + 2\mu + a_m(r_A + \mu)(r_B + \mu) + \delta, \\
0 &< r_A + r_B + \mu + a_m(r_A + \mu)(r_B + \mu) + \delta, \qquad \text{(SI1.12)}
\end{aligned}$$

because from eq. (SI1.6), we know that $W_i = a_m(r_i + \mu)$.

The only biologically relevant scenario is one in which the population does not go extinct. This requires non-negative intrinsic growth rates, i.e. $r_i \geq 0$. From eq. (SI1.12), it follows that when $r_i \geq 0$, the inequality is always satisfied because all other terms in the right-hand side are strictly positive ($a_m > 0$ and $\delta > 0$).

> As a consequence, when selection acts on fecundity, $\frac{\partial c_f}{\partial a_m} < 0$, indicating that the selection coefficient always decreases with increasing age at maturity in a viable population (i.e. a population with non-negative intrinsic growth rate).

#### 2 Scenario. Fitness advantage in pre-maturity mortality

In the second scenario, we consider the case in which genotype A experiences lower pre-maturity mortality, such that $\delta_A < \delta_B$, whereas all other demographic rates are equal. Hence, $\beta_A = \beta_B = \beta$ and $\mu_A = \mu_B = \mu$. In such a case, the selection coefficient is

$$c_\delta = \left(\frac{W_A}{a_m} - \mu\right) - \left(\frac{W_B}{a_m} - \mu\right) = \frac{1}{a_m}(W_A - W_B), \qquad \text{(SI1.13)}$$

where $W_A = W\big(a_m \beta e^{a_m(\mu - \delta_A)}\big)$ and $W_B = W\big(a_m \beta e^{a_m(\mu - \delta_B)}\big)$. To understand how $c_\delta$ depends on $a_m$, we calculate the derivative of this equation with respect to $a_m$

$$\frac{\partial c_\delta}{\partial a_m} = -\frac{1}{a_m^2}(W_A - W_B) + \frac{1}{a_m}\left(\frac{\partial W_A}{\partial a_m} - \frac{\partial W_B}{\partial a_m}\right). \qquad \text{(SI1.14)}$$

Using the definition of the derivative of the Lambert W function given by eq. (SI1.10), we obtain

$$\frac{\partial W_i}{\partial a_m} = \frac{W_i}{z_i(1 + W_i)} \frac{\partial z_i}{\partial a_m}.$$

where $z_i(a_m) = a_m \, \beta \, e^{a_m(\mu - \delta_i)}$, and

$$\frac{\partial z_i}{\partial a_m} = \beta e^{a_m(\mu - \delta_i)}\big[1 + (\mu - \delta_i)a_m\big] = \frac{z_i}{a_m}\big[1 + (\mu - \delta_i)a_m\big].$$

Therefore,

$$\frac{\partial W_i}{\partial a_m} = \frac{W_i}{1+W_i}\,\frac{1+(\mu-\delta_i)a_m}{a_m}.$$

Substituting into the equation (SI1.14), it follows that

$$\begin{aligned}
\frac{\partial c_\delta}{\partial a_m} &= \frac{1}{a_m^2}\left[-\big(W_A-W_B\big)+\left(\frac{W_A}{1+W_A}\left(1+(\mu-\delta_A)a_m\right)-\frac{W_B}{1+W_B}\left(1+(\mu-\delta_B)a_m\right)\right)\right]\\
&= \frac{1}{a_m^2}\left[\frac{W_A\left(1+(\mu-\delta_A)a_m\right)-W_A(1+W_A)}{1+W_A}-\frac{W_B\left(1+(\mu-\delta_B)a_m\right)-W_B(1+W_B)}{1+W_B}\right]\\
&= \frac{1}{a_m^2}\left[\frac{W_A\left((\mu-\delta_A)a_m-W_A\right)}{1+W_A}-\frac{W_B\left((\mu-\delta_B)a_m-W_B\right)}{1+W_B}\right]\\
&= \frac{1}{a_m^2}\left[\frac{W_A\left((\mu-\delta_A)a_m-(r_A+\mu)a_m\right)}{1+W_A}-\frac{W_B\left((\mu-\delta_B)a_m-(r_B+\mu)a_m\right)}{1+W_B}\right]\\
&= \frac{1}{a_m^2}\left[\frac{-W_A\left((r_A+\delta_A)a_m\right)}{1+W_A}+\frac{W_B\left((r_B+\delta_B)a_m\right)}{1+W_B}\right]\\
&= \frac{1}{a_m}\left[\frac{-a_m(r_A+\mu)\left(r_A+\delta_A\right)}{1+a_m(r_A+\mu)}+\frac{a_m(r_B+\mu)\left(r_B+\delta_B\right)}{1+a_m(r_B+\mu)}\right]\\
&= -\frac{(r_A+\mu)\left(r_A+\delta_A\right)}{1+a_m(r_A+\mu)}+\frac{(r_B+\mu)\left(r_B+\delta_B\right)}{1+a_m(r_B+\mu)}\\
&= \frac{-(1+a_m(r_B+\mu))(r_A+\mu)\left(r_A+\delta_A\right)+(1+a_m(r_A+\mu))(r_B+\mu)\left(r_B+\delta_B\right)}{(1+a_m(r_A+\mu))(1+a_m(r_B+\mu))} \qquad \text{(SI1.15)}
\end{aligned}$$

From step three to four and five to six, we replace $W_i$ using the identity $W_i = a_m(r_i+\mu)$ from eq. (SI1.6). Because the denominator has only non-negative terms ($a_m > 0$, $\mu > 0$, and $r_i \geq 0$ to enable population persistence), it is strictly positive. Therefore, the sign of $\frac{\partial c_\delta}{\partial a_m}$ is determined by the numerator, $N_\delta$.

$$\begin{aligned}
N_\delta &= -(1+a_m(r_B+\mu))(r_A+\mu)\left(r_A+\delta_A\right)+(1+a_m(r_A+\mu))(r_B+\mu)\left(r_B+\delta_B\right)\\
&= (r_B+\mu)\left(r_B+\delta_B\right)-(r_A+\mu)\left(r_A+\delta_A\right)+a_m\left[(r_A+\mu)(r_B+\mu)(r_B+\delta_B-r_A-\delta_A)\right]\\
&= (\delta_B-\delta_A)[\mu+r_A+a_m(r_A+\mu)(r_B+\mu)]-(r_A-r_B)[(r_A+r_B)+\mu+\delta_B+a_m(r_A+\mu)(r_B+\mu)]\\
&= (\delta_B-\delta_A)[(r_A+\mu)(1+a_m(r_B+\mu))]-(r_A-r_B)[(r_A+\mu)\big(1+a_m(r_B+\mu)\big)+(r_B+\delta_B)]
\end{aligned} \qquad \text{(SI1.16)}$$

Hence,

$$N_\delta > 0 \iff \frac{P}{P+(r_B+\delta_B)} > \frac{(r_A-r_B)}{(\delta_B-\delta_A)}, \qquad \text{(SI1.17)}$$

where $P = (r_A+\mu)(1+a_m(r_B+\mu))$.

By the mean value theorem, for some $\xi \in (\delta_A, \delta_B)$,

$$r_A - r_B \;=\; \left(\frac{\partial r}{\partial \delta}\right)_{\delta=\xi}(\delta_A-\delta_B) \;=\; (\delta_B-\delta_A)\frac{a_m\left(r(\xi)+\mu\right)}{1+a_m\left(r(\xi)+\mu\right)}$$

because from the implicit equation $\beta = (r+\mu)\,e^{(r+\delta)a_m}$ (eq.SI1.2), we calculate the derivative

$$\frac{\partial r}{\partial \delta} = -\frac{a_m\,(r+\mu)}{1+a_m\,(r+\mu)}.$$

Thus, $(r_A-r_B)/(\delta_B-\delta_A) = x/(1+x)$ with $x = a_m\big(r(\xi)+\mu\big) \in [a_m(r_B+\mu),\, a_m(r_A+\mu)]$, which we can replace in eq. (SI1.17), yielding

$$N_\delta > 0 \iff \frac{P}{P+(r_B+\delta_B)} > \frac{x}{1+x}, \qquad x \in \big[a_m(r_B+\mu),\, a_m(r_A+\mu)\big].$$

Equivalently,

$$N_\delta > 0 \iff P > x(r_B + \delta_B), \qquad x \in \left[a_m(r_B + \mu),\, a_m(r_A + \mu)\right].$$

Because $a_m(r_A + \mu) > a_m(r_B + \mu)$, a sufficient condition for $N_\delta > 0$ is

$$\begin{aligned} P &> a_m(r_A + \mu)(r_B + \delta_B) \\ (r_A + \mu)(1 + a_m(r_B + \mu)) &> a_m(r_A + \mu)(r_B + \delta_B) \\ 1 &> a_m(r_B + \delta_B - r_B - \mu) \\ 1 &> a_m(\delta_B - \mu) \end{aligned} \tag{SI1.18}$$

This condition is always satisfied when $\mu > \delta_B$, thus $\frac{\partial c_\delta}{\partial a_m} > 0$ in such a case. Conversely, when $\mu < \delta_B$, the condition is satisfied only if $a_m < 1/(\delta_B - \mu)$, otherwise, when $a_m$ surpass the threshold $1/(\delta_B - \mu)$, the condition is not longer satisfied, and thus $\frac{\partial c_\delta}{\partial a_m} < 0$.

In the following, we explore the asymptotic behavior of $N_\delta$ to gain insight into the behavior of the selection coefficient in the limits $a_m \to 0$ and $a_m \to \infty$.

**Asymptotic behavior: very small age at maturity** We begin by studying the case when $a_m$ is very small, in the limit $a_m \to 0$. In this case:

$$W_i = W(z_i) \approx z_i$$

Therefore,

$$r_i \approx \frac{z_i}{a_m} - \mu = \frac{a_m\, \beta\, e^{a_m(\mu - \delta_A)}}{a_m} - \mu = \beta e^{a_m(\mu - \delta_A)} - \mu,$$

which approximates as

$$\lim_{a_m \to 0} r_i \approx \beta - \mu.$$

This approximation is consistent with models that consider only birth and death processes, and exclude maturation, which define the growth rate as the difference between the two processes. Using this approximation and the definition of $N_\delta$, we have

$$\begin{aligned} \lim_{a_m \to 0} N_\delta &\approx r_B^2 - r_A^2 + \mu[r_B - r_A + \delta_B - \delta_A] + r_B\delta_B - r_A\delta_A \\ &\approx \mu(\delta_B - \delta_A) + (\beta - \mu)\delta_B - (\beta - \mu)\delta_A \\ &\approx \mu(\delta_B - \delta_A) + (\beta - \mu)(\delta_B - \delta_A) \end{aligned} \tag{SI1.19}$$

To satisfy $r_i \geq 0$ (necessary condition for population persistence), $\beta > \mu$. Additionally, we know that $\delta_B > \delta_A$, therefore $\lim_{a_m \to 0} N_\delta > 0$. Consequently, when selection acts on pre-maturity survival and the age at maturity is small, $\frac{\partial c_\delta}{\partial a_m} > 0$ and thus the selection coefficient increases with age at maturity.

**Asymptotic behavior: very large age at maturity** We continue the analysis considering now the case when $a_m$ is very large, in the limit $a_m \to \infty$. In such case, a population is in decline because to persist, a population requires a non–negative intrinsic growth rate $r_i \geq 0$. Using eq.(SI1.6), this condition imposes an upper limit on the age at maturity,

$$a_m \leq a_m^{\max} = \frac{1}{\delta_i} \ln\left(\frac{\beta_i}{\mu_i}\right). \tag{SI1.20}$$

Bearing this in mind, we proceed with the analysis. The behavior of the argument of the Lambert W function, $z_i = a_m\, \beta\, e^{a_m(\mu - \delta_i)}$ depends on the sign of $(\mu - \delta_i)$. If $\mu > \delta_i$,

$$\lim_{a_m \to \infty} z_i \approx \infty.$$

Whereas, if $\mu < \delta_i$,

$$\lim_{a_m \to \infty} z_i \approx 0.$$

We therefore examine the two subcases separately. In the first subcase, when post-maturity mortality is higher than pre-maturity mortality, $\mu > \delta_i$, in the limit $a_m \to \infty$, we have that

$$W_i = W(z_i) \approx \ln z_i - \ln(\ln z_i),$$

where $\ln z_i = \ln a_m + \ln \beta + a_m(\mu - \delta_i)$, which can be approximated as

$$\lim_{a_m \to \infty} \ln z_i \approx a_m(\mu - \delta_i).$$

Hence,

$$\lim_{a_m \to \infty} \ln(\ln z_i) \approx \ln(a_m(\mu - \delta_i)) = \ln a_m + \ln(\mu - \delta_i).$$

Therefore, using the definition $r_i = \frac{W_i}{a_m} - \mu$, we obtain the approximation:

$$\begin{aligned}
\lim_{a_m \to \infty} r_i &\approx \frac{\ln z_i - \ln(\ln z_i)}{a_m} - \mu \\
&\approx \frac{\ln a_m + \ln \beta + a_m(\mu - \delta_i) - \ln a_m - \ln(\mu - \delta_i)}{a_m} - \mu \\
&\approx \frac{\ln \beta}{a_m} + (\mu - \delta_i) - \frac{\ln(\mu - \delta_i)}{a_m} - \mu \\
&\approx -\delta_i + \underbrace{\frac{1}{a_m}\left(\ln \beta - \ln(\mu - \delta_i)\right)}_{O(\frac{1}{a_m})}
\end{aligned} \tag{SI1.21}$$

Note that in this limit $r_i \approx -\delta_i$, so the population is declining, as mentioned above. We use eq. (SI1.21) to approximate $N_\delta$:

$$\begin{aligned}
\lim_{a_m \to \infty} N_\delta &\approx a_m(r_A + \mu)(r_B + \mu)(r_B - r_A + \delta_B - \delta_A) \\
&\approx a_m(\delta_A + \mu)(\delta_B + \mu)(\delta_B - \delta_A + \delta_B - \delta_A)
\end{aligned}$$

Since all the terms in this expression are strictly positive ($a_m > 0$, $\delta_A > 0$, $\delta_B > 0$, and $\mu > 0$) and $\delta_B > \delta_A$, $\lim_{a_m \to 0} N_\delta > 0$. Consequently, when selection acts on pre-maturity mortality, and this is smaller than post-maturity mortality, $\frac{\partial c_\delta}{\partial a_m} > 0$ and thus the selection coefficient increases with age at maturity.

Now, we examine the second subcase, in which pre-maturity mortality is higher than post-maturity mortality ($\mu_i < \delta$). Because in this limit the argument of the Lambert W function approaches zero ($\lim_{a_m \to \infty} z_i \approx 0$), we use the following approximation

$$W_i = W(z_i) \approx z_i = a_m \beta \, e^{a_m(\mu - \delta_i)},$$

which yields

$$r_i \approx \frac{a_m \beta \, e^{a_m(\mu - \delta_i)}}{a_m} - \mu = \beta \, e^{a_m(\mu - \delta_i)} - \mu.$$

Let $\beta \, e^{a_m(\mu - \delta_i)} = \varepsilon_i$. Hence,

$$r_i + \mu \approx \varepsilon_i \to 0^+, \qquad r_B - r_A \approx \varepsilon_B - \varepsilon_A.$$

Substituting these limits into the terms of eq. (SI1.16), which we analyze separately, we have

$$\begin{aligned}
r_B^2 - r_A^2 =& (r_B - r_A)(r_B + r_A) \\
&\approx (\varepsilon_B - \varepsilon_A)(\varepsilon_A + \varepsilon_B - 2\mu) \\
&\approx -2\mu(\varepsilon_B - \varepsilon_A) + \varepsilon_B^2 - \varepsilon_A^2 \\
\mu\,[r_B - r_A + \delta_B - \delta_A] &\approx \mu(\varepsilon_B - \varepsilon_A) + \mu(\delta_B - \delta_A), \\
r_B\delta_B - r_A\delta_A &\approx \delta_B(\varepsilon_B - \mu) - \delta_A(\varepsilon_A - \mu) \\
&\approx \delta_B\varepsilon_B - \delta_B\mu - \delta_A\varepsilon_A + \delta_A\mu \\
&\approx -\mu(\delta_B - \delta_A) + \delta_B\varepsilon_B - \delta_A\varepsilon_A
\end{aligned}$$

and

$$a_m(r_A + \mu)(r_B + \mu)(r_B - r_A + \delta_B - \delta_A) \approx a_m\,\varepsilon_A\varepsilon_B(\varepsilon_B - \varepsilon_A + \delta_B - \delta_A).$$

Collecting,

$$\begin{aligned}
N_\delta =& -2\mu(\varepsilon_B - \varepsilon_A) + \varepsilon_B^2 - \varepsilon_A^2 + \mu(\varepsilon_B - \varepsilon_A) + \mu(\delta_B - \delta_A) - \mu(\delta_B - \delta_A) + \delta_B\varepsilon_B - \delta_A\varepsilon_A \\
&+ a_m\,\varepsilon_A\varepsilon_B(\varepsilon_B - \varepsilon_A + \delta_B - \delta_A) \\
=& -\mu\varepsilon_B + \mu\varepsilon_A + \varepsilon_B\delta_B - \varepsilon_A\delta_A + \varepsilon_B^2 - \varepsilon_A^2 + a_m\,\varepsilon_A\varepsilon_B(\varepsilon_B - \varepsilon_A + \mu_A - \mu_B) \\
=& \varepsilon_B(\delta_B - \mu) - \varepsilon_A(\delta_A - \mu) + \underbrace{\varepsilon_B^2 - \varepsilon_A^2}_{O(\varepsilon^2)} + \underbrace{a_m\,\varepsilon_A\varepsilon_B(\delta_B - \delta_A)}_{O(a_m\varepsilon^2)} + \underbrace{a_m\,\varepsilon_A\varepsilon_B(\varepsilon_B - \varepsilon_A)}_{O(a_m\varepsilon^3)}
\end{aligned}$$

Since $\delta_A < \delta_B$, in the limit $a_m \to \infty$, $\varepsilon_B$ is exponentially smaller than $\varepsilon_A$ because

$$\frac{\varepsilon_B}{\varepsilon_A} = e^{a_m(\delta_A - \delta_B)} \to 0 \qquad \text{as } a_m \to \infty.$$

Therefore, we have

$$0 < \varepsilon_B \ll \varepsilon_A \ll 1.$$

Consequently,

$$\lim_{a_m \to \infty} N_\delta \approx \varepsilon_B(\delta_B - \mu) - \varepsilon_A(\delta_A - \mu) + O(\varepsilon^2) + O(a_m\varepsilon^2) + O(a_m\varepsilon^3) > 0$$

because $\delta_A - \mu > 0$ and $\varepsilon_A \gg \varepsilon_B$. Thus, $\lim_{a_m \to \infty} \frac{\partial c_\delta}{\partial a_m} = 0$, yet for large but finite $a_m$, $\frac{\partial c_\delta}{\partial a_m} < 0$ when $\mu < \delta_i$.

The analysis of the asymptotic behavior is thus consistent with our analytical findings, which we summarize as:

When selection acts on pre-maturity mortality, two regimes occur:

1. when immature individuals experience a higher mortality rate than mature ones, $\mu < \delta$, the most common case in nature, the selection coefficient initially increases with age at maturity until the threshold $a_m = 1/(\delta - \mu)$, and then decreases. Therefore, the smaller the difference between immature and mature mortality rates, the larger the range over which the selection coefficient increases with age at maturity.

2. when mature individuals experience a higher mortality rate than immature ones, $\mu > \delta$, the selection coefficient always increases monotonically with age at maturity.

#### 3 Scenario. Fitness advantage in post-maturity mortality

In the third scenario, we consider the case in which genotype A experiences lower post-maturity mortality, such that $\mu_A < \mu_B$, whereas all other demographic rates are equal. Hence, $\beta_A = \beta_B = \beta$ and $\delta_A = \delta_B = \delta$. In such a case, the selection coefficient is

$$c_\mu = \left(\frac{W_A}{a_m} - \mu_A\right) - \left(\frac{W_B}{a_m} - \mu_B\right) = \frac{1}{a_m}\left(W_A - W_B\right) - (\mu_A - \mu_B), \tag{SI1.22}$$

where $W_A = W\big(a_m\beta e^{a_m(\mu_A-\delta)}\big)$ and $W_B = W\big(a_m\beta e^{a_m(\mu_B-\delta)}\big)$. To understand how $c_\mu$ depends on $a_m$, we calculate the derivative of this equation with respect to $a_m$

$$\frac{\partial c_\mu}{\partial a_m} = -\frac{1}{a_m^2}\left(W_A - W_B\right) + \frac{1}{a_m}\left(\frac{\partial W_A}{\partial a_m} - \frac{\partial W_B}{\partial a_m}\right). \tag{SI1.23}$$

Using the definition of the derivative of the Lambert W function given by eq. (SI1.10), we obtain

$$\frac{\partial W_i}{\partial a_m} = \frac{W_i}{z_i(1+W_i)}\frac{\partial z_i}{\partial a_m}.$$

where $z_i(a_m) = a_m\,\beta\, e^{a_m(\mu_i-\delta)}$, and

$$\frac{\partial z_i}{\partial a_m} = \beta e^{a_m(\mu_i-\delta)}\big[1 + (\mu_i - \delta)a_m\big] = \frac{z_i}{a_m}\big[1 + (\mu_i - \delta)a_m\big].$$

Therefore,

$$\frac{\partial W_i}{\partial a_m} = \frac{W_i}{1+W_i}\,\frac{1 + (\mu_i - \delta)a_m}{a_m}.$$

Substituting into the equation (SI1.23), it follows that

$$\begin{aligned}
\frac{\partial c_\mu}{\partial a_m} &= \frac{1}{a_m^2}\left[-\big(W_A - W_B\big) + \left(\frac{W_A}{1+W_A}\left(1 + (\mu_A-\delta)a_m\right) - \frac{W_B}{1+W_B}\left(1 + (\mu_B-\delta)a_m\right)\right)\right] \\
&= \frac{1}{a_m^2}\left[\frac{W_A\left(1 + (\mu_A-\delta)a_m\right) - W_A(1+W_A)}{1+W_A} - \frac{W_B\left(1 + (\mu_B-\delta)a_m\right) - W_B(1+W_B)}{1+W_B}\right] \\
&= \frac{1}{a_m^2}\left[\frac{W_A\left((\mu_A-\delta)a_m - W_A\right)}{1+W_A} - \frac{W_B\left((\mu_B-\delta)a_m - W_B\right)}{1+W_B}\right] \\
&= \frac{1}{a_m^2}\left[\frac{W_A\left((\mu_A-\delta)a_m - (r_A+\mu_A)a_m\right)}{1+W_A} - \frac{W_B\left((\mu_B-\delta)a_m - (r_B+\mu_B)a_m\right)}{1+W_B}\right] \\
&= \frac{1}{a_m^2}\left[\frac{-W_A\left((r_A+\delta)a_m\right)}{1+W_A} + \frac{W_B\left((r_B+\delta)a_m\right)}{1+W_B}\right] \\
&= \frac{1}{a_m}\left[\frac{-a_m(r_A+\mu_A)\,(r_A+\delta)}{1+a_m(r_A+\mu_A)} + \frac{a_m(r_B+\mu_B)\,(r_B+\delta)}{1+a_m(r_B+\mu_B)}\right] \\
&= -\frac{(r_A+\mu_A)\,(r_A+\delta)}{1+a_m(r_A+\mu_A)} + \frac{(r_B+\mu_B)\,(r_B+\delta)}{1+a_m(r_B+\mu_B)} \\
&= \frac{-(1+a_m(r_B+\mu_B))(r_A+\mu_A)\,(r_A+\delta) + (1+a_m(r_A+\mu_A))(r_B+\mu_B)\,(r_B+\delta)}{(1+a_m(r_A+\mu_A))(1+a_m(r_B+\mu_B))}
\end{aligned} \tag{SI1.24}$$

From step three to four and five to six, we replace $W_i$ using the identity $W_i = a_m(r_i + \mu_i)$ from eq. (SI1.6). Because the denominator has only non-negative terms ($a_m > 0$, $\mu_i > 0$, and $r_i \geq 0$ to enable population persistence), it is strictly positive. Therefore, the sign of $\frac{\partial c_\mu}{\partial a_m}$ is determined by the numerator, $N$.

$$
\begin{aligned}
N_\mu = & -(1+a_m(r_B+\mu_B))(r_A+\mu_A)\,(r_A+\delta)+(1+a_m(r_A+\mu_A))(r_B+\mu_B)\,(r_B+\delta)\\
=&(r_B+\mu_B)\,(r_B+\delta)-(r_A+\mu_A)\,(r_A+\delta)+a_m\left[(r_A+\mu_A)(r_B+\mu_B)(r_B+\delta-r_A-\delta)\right]\\
=&r_B^2-r_A^2+\delta[r_B-r_A+\mu_B-\mu_A]+r_B\mu_B-r_A\mu_A+a_m\left[(r_A+\mu_A)(r_B+\mu_B)(r_B-r_A)\right]\\
=&(r_B-r_A)(r_B+r_A)+\delta(r_B-r_A)+\delta(\mu_B-\mu_A)+\mu_A(r_B-r_A)+r_B(\mu_B-\mu_A)\\
&+a_m\left[(r_A+\mu_A)(r_B+\mu_B)(r_B-r_A)\right]\\
=&(r_B-r_A)[r_A+r_B+\delta+\mu_A+a_m(r_A+\mu_A)(r_B+\mu_B)]+(\mu_B-\mu_A)(\delta+r_B)\\
=&-(r_A-r_B)[\delta+r_B+(r_A+\mu_A)[1+a_m(r_B+\mu_B)]+(\mu_B-\mu_A)(\delta+r_B)
\end{aligned}
\tag{SI1.25}
$$

Hence,

$$
N_\mu<0 \iff \frac{\delta+r_B}{\delta+r_B+(r_A+\mu_A)[1+a_m(r_B+\mu_B)]}<\frac{(r_A-r_B)}{(\mu_B-\mu_A)}. \tag{SI1.26}
$$

By the mean value theorem, for some $\xi \in (\mu_A, \mu_B)$,

$$
r_A-r_B \;=\; \left(\frac{\partial r}{\partial \mu}\right)_{\mu=\xi}(\mu_A-\mu_B) \;=\; (\mu_B-\mu_A)\frac{1}{1+a_m\left(r(\xi)+\mu(\xi)\right)} \tag{SI1.27}
$$

because from the implicit equation $\beta=(r+\mu)\,e^{(r+\delta)a_m}$ (eq.SI1.2), we calculate the derivative

$$
\frac{\partial r}{\partial \mu}=-\frac{1}{1+a_m\,(r+\mu)}.
$$

Thus, $(r_A-r_B)/(\mu_B-\mu_A)=1/(1+x)$ with $x=a_m\big(r(\xi)+\mu(\xi)\big)\in[a_m(r_B+\mu_B),\, a_m(r_A+\mu_A)]$, which we can replace in eq. (SI1.26), yielding

$$
N_\mu<0 \iff \frac{\delta+r_B}{\delta+r_B+(r_A+\mu_A)[1+a_m(r_B+\mu_B)]}<\frac{1}{1+x}, \qquad x\in\left[a_m(r_B+\mu_B),\, a_m(r_A+\mu_A)\right].
$$

We define $g(\mu)=r(\mu)+\mu$, which is an increasing function because

$$
g'(\mu) \;=\; \frac{\partial r}{\partial \mu}+1 \;=\; 1-\frac{1}{1+a_m(r+\mu)} \;=\; \frac{a_m(r+\mu)}{1+a_m(r+\mu)} \;>\; 0,
$$

so

$$
\frac{1}{1+a_m(r_B+\mu_B)} \;<\; \frac{r_A-r_B}{\mu_B-\mu_A} \;<\; \frac{1}{1+a_m(r_A+\mu_A)}. \tag{SI1.28}
$$

Therefore, a sufficient condition for $N_\mu<0$ is

$$
\frac{\delta+r_B}{\delta+r_B+(r_A+\mu_A)[1+a_m(r_B+\mu_B)]} \;<\; \frac{1}{1+a_m(r_B+\mu_B)}.
$$

Equivalently,

$$
\begin{aligned}
(\delta+r_B)\left(1+a_m(r_B+\mu_B)\right) \;&<\; \delta+r_B+(r_A+\mu_A)[1+a_m(r_B+\mu_B)]\\
\delta+r_B+a_m(r_B+\mu_B)(\delta+r_B) \;&<\; \delta+r_B+(r_A+\mu_A)+a_m(r_B+\mu_B)(r_A+\mu_A)\\
0 \;&<\; (r_A+\mu_A)+a_m(r_B+\mu_B)(r_A-r_B+\mu_A-\delta)\\
\frac{-(r_A+\mu_A)}{(r_B+\mu_B)(r_A-r_B+\mu_A-\delta)} \;&<a_m.
\end{aligned}
\tag{SI1.29}
$$

Because $r_A-r_B>0$, the left-hand side is negative, and thus the inequality is satisfied, whenever $\mu_A>\delta$ in a scenario of population persistence ($r_i\geq 0$). Hence, when post-maturity mortality is higher than pre-maturity mortality, $N_\mu < 0$, and thus $\frac{\partial c_\mu}{\partial a_m} < 0$, indicating that the selection coefficient decreases with age at maturity.

Yet, from this condition, it is difficult to understand how $N_\mu$ behaves when $\mu_A < \mu_B < \delta$. Thus, we now explore

$$N_\mu > 0 \iff \frac{\delta + r_B}{\delta + r_B + (r_A + \mu_A)[1 + a_m(r_B + \mu_B)]} > \frac{(r_A - r_B)}{(\mu_B - \mu_A)}. \tag{SI1.30}$$

Replacing $(r_A - r_B)/(\mu_B - \mu_A) = 1/(1+x)$ with $x = a_m\big(r(\xi) + \mu(\xi)\big) \in [a_m(r_B + \mu_B),\, a_m(r_A + \mu_A)]$ in eq. (SI1.30), yields

$$N_\mu > 0 \iff \frac{\delta + r_B}{\delta + r_B + (r_A + \mu_A)[1 + a_m(r_B + \mu_B)]} > \frac{1}{1+x}, \qquad x \in \big[a_m(r_B + \mu_B),\, a_m(r_A + \mu_A)\big].$$

From eq. SI1.28, we know that a sufficient condition for $N_\mu > 0$ is

$$\frac{\delta + r_B}{\delta + r_B + (r_A + \mu_A)[1 + a_m(r_B + \mu_B)]} > \frac{1}{1 + a_m(r_A + \mu_A)}.$$

Equivalently,

$$\begin{aligned}
(\delta + r_B)\big(1 + a_m(r_A + \mu_A)\big) &> \delta + r_B + (r_A + \mu_A)[1 + a_m(r_B + \mu_B)] \\
\delta + r_B + a_m(r_A + \mu_A)(\delta + r_B) &> \delta + r_B + (r_A + \mu_A)[1 + a_m(r_B + \mu_B)] \\
a_m(\delta + r_B) &> 1 + a_m(r_B + \mu_B) \\
a_m &> \frac{1}{(\delta - \mu_B)}
\end{aligned} \tag{SI1.31}$$

Therefore, $N_\mu > 0$ when $a_m > 1/(\delta - \mu_B)$, provided that $\delta > \mu_B$. Hence, when pre-maturity mortality is higher than pre-maturity mortality, the selection coefficient first decreases until the age at maturity reaches $1/(\delta - \mu_B)$, and then increases with age at maturity.

In the following, we explore the asymptotic behavior of $N_\mu$ to gain insight into the behavior of the selection coefficient in the limits $a_m \to 0$ and $a_m \to \infty$.

**Asymptotic behavior: Very small age at maturity** We begin by studying the case when $a_m$ is very small, in the limit $a_m \to 0$. In this case:

$$W_i = W(z_i) \approx z_i$$

Therefore,

$$r_i = \frac{W(z_i)}{a_m} - \mu_i \approx \frac{a_m\, \beta\, e^{a_m(\mu - \delta_A)}}{a_m} - \mu_i,$$

which approximates as

$$\lim_{a_m \to 0} r_i \approx \beta - \mu_i.$$

We then use this to approximate $N_\mu$:

$$\begin{aligned}
\lim_{a_m \to 0} N_\mu \approx& (\beta - \mu_B)^2 - (\beta - \mu_A)^2 + \delta[(\beta - \mu_B) - (\beta - \mu_A) + \mu_B - \mu_A] + (\beta - \mu_B)\mu_B - (\beta - \mu_A)\mu_A \\
&+ a_m\left[(\beta - \mu_A) + \mu_A)((\beta - \mu_B) + \mu_B)((\beta - \mu_B) - (\beta - \mu_A))\right] \\
\approx& (\beta^2 - 2\beta\mu_B + \mu_B^2) - (\beta^2 + 2\beta\mu_A + \mu_A^2) + \beta\mu_B - \mu_B^2 - \beta\mu_A + \mu_A^2 + a_m\beta^2(\mu_A - \mu_B) \\
\approx& \beta(\mu_A - \mu_B) + a_m\beta^2(\mu_A - \mu_B)
\end{aligned}$$

Since $\mu_A < \mu_B$, $\lim_{a_m \to 0} N_\mu < 0$. Consequently, when selection acts on post-maturity survival and age at maturity is small, $\frac{\partial c_\mu}{\partial a_m} < 0$ and thus the selection coefficient decreases with age at maturity.

**Asymptotic behavior: Very large age at maturity** We now consider the case when $a_m$ is very large, in the limit $a_m \to \infty$. In such case, the behavior of the argument of the Lambert W function, $z_i = a_m\,\beta\, e^{a_m(\mu_i-\delta)}$ depends on the sign of $(\mu_i - \delta)$. If $\mu_i > \delta$,

$$\lim_{a_m \to \infty} z_i \approx \infty.$$

Whereas, if $\mu_i < \delta$,

$$\lim_{a_m \to \infty} z_i \approx 0.$$

We therefore examine the two subcases separately. In the first subcase, when post-maturity mortality is higher than pre-maturity mortality, $\mu_i > \delta$, in the limit $a_m \to \infty$, we have that

$$W_i = W(z_i) \approx \ln z_i - \ln(\ln z_i),$$

where $\ln z_i = \ln a_m + \ln\beta + a_m(\mu_i - \delta)$, which can be approximated as

$$\lim_{a_m \to \infty} \ln z_i \approx a_m(\mu_i - \delta).$$

Hence,

$$\lim_{a_m \to \infty} \ln(\ln z_i) \approx \ln(a_m(\mu_i - \delta)) = \ln a_m + \ln(\mu_i - \delta).$$

Therefore, using the definition $r_i = \frac{W_i}{a_m} - \mu$, we obtain the approximation:

$$\begin{aligned} \lim_{a_m \to \infty} r_i &\approx \frac{\ln z_i - \ln(\ln z_i)}{a_m} - \mu \\ &\approx \frac{\ln a_m + \ln\beta + a_m(\mu_i - \delta) - \ln a_m - \ln(\mu_i - \delta)}{a_m} - \mu_i \\ &\approx -\delta + \frac{1}{a_m}\,\ln\frac{\beta}{\mu_i - \delta}. \end{aligned} \tag{SI1.32}$$

Hence, in the limit $a_m \to \infty$,

$$r_B - r_A \approx \frac{1}{a_m}\left[\ln\frac{\beta}{\mu_B - \delta} - \ln\frac{\beta}{\mu_A - \delta}\right] = \frac{1}{a_m}\,\ln\frac{\mu_A - \delta}{\mu_B - \delta} \;<\; 0, \qquad r_B + r_A \approx -2\delta.$$

Using this approximation for the terms in $N_\mu$:

$$\begin{aligned} r_B^2 - r_A^2 = (r_B + r_A)(r_B - r_A) \;&\approx\; \frac{-2\delta}{a_m}\,\ln\frac{\mu_A - \delta}{\mu_B - \delta}, \\ \delta\left[r_B - r_A + \mu_B - \mu_A\right] + r_B\mu_B - r_A\mu_A =& \delta\,(r_B - r_A) + \delta(\mu_B - \mu_A) - \delta(\mu_B - \mu_A) \\ &+ \mu_B(r_B + \delta) - \mu_A(r_A + \delta) \\ =& \delta\,(r_B - r_A) + \mu_B(r_B + \delta) - \mu_A(r_A + \delta) \\ \approx& \frac{\delta}{a_m}\,\ln\frac{\mu_A - \delta}{\mu_B - \delta} \;+\; \frac{1}{a_m}\left[\mu_B\ln\frac{\beta}{\mu_B - \delta} - \mu_A\ln\frac{\beta}{\mu_A - \delta}\right], \\ \approx& \frac{1}{a_m}\left[\delta\ln\frac{\mu_A - \delta}{\mu_B - \delta} \;+\; \mu_B\ln\frac{\beta}{\mu_B - \delta} - \mu_A\ln\frac{\beta}{\mu_A - \delta}\right], \\ a_m(r_A + \mu_A)(r_B + \mu_B)(r_B - r_A) =& a_m(r_B - r_A)(r_A + \mu_A)(r_B + \mu_B) \\ \approx& \ln\frac{\mu_A - \delta}{\mu_B - \delta}\left(\tfrac{1}{a_m}\ln\tfrac{\beta}{\mu_A - \delta} - \delta + \mu_A\right)\left(\tfrac{1}{a_m}\ln\tfrac{\beta}{\mu_B - \delta} - \delta + \mu_B\right) \end{aligned}$$

Collecting,

$$N_\mu = \ln\frac{\mu_A-\delta}{\mu_B-\delta}\Big(\underbrace{\tfrac{1}{a_m}\ln\tfrac{\beta}{\mu_A-\delta}}_{O(\frac{1}{a_m})}-\delta+\mu_A\Big)\Big(\underbrace{\tfrac{1}{a_m}\ln\tfrac{\beta}{\mu_B-\delta}}_{O(\frac{1}{a_m})}-\delta+\mu_B\Big)$$
$$+\underbrace{\frac{1}{a_m}\Big[-2\delta\,\ln\frac{\mu_A-\delta}{\mu_B-\delta}+\delta\ln\frac{\mu_A-\delta}{\mu_B-\delta}+\mu_B\ln\frac{\beta}{\mu_B-\delta}-\mu_A\ln\frac{\beta}{\mu_A-\delta}\Big]}_{O(\frac{1}{a_m})}$$

Therefore,

$$\lim_{a_m\to\infty} N_\mu \;=\; \ln\frac{\mu_A-\delta}{\mu_B-\delta}(\mu_A-\delta)(\mu_B-\delta). \tag{SI1.33}$$

Because $\mu_A-\delta<\mu_B-\delta \Rightarrow \ln(\frac{\mu_A-\delta}{\mu_B-\delta})<0$. Therefore, $\lim_{a_m\to\infty} N_\mu<0$ and thus $\frac{\partial c_\mu}{\partial a_m}<0$ in the subcase $\mu_i>\delta$.

Now, we examine the second subcase, in which pre-maturity mortality is higher than post-maturity mortality ($\mu_i<\delta$). Because in this limit the argument of the Lambert W function approaches zero ($\lim_{a_m\to\infty} z_i\approx 0$), we use the following approximation

$$W_i=W(z_i)\approx z_i=a_m\,\beta\,e^{a_m(\mu_i-\delta)},$$

which yields

$$r_i\approx\frac{a_m\beta\,e^{a_m(\mu_i-\delta)}}{a_m}-\mu_i=\beta\,e^{a_m(\mu_i-\delta)}-\mu_i.$$

Let $\beta\,e^{a_m(\mu_i-\delta)}=\varepsilon_i$. Hence,

$$r_i+\mu_i\approx\varepsilon_i\to 0^+,\qquad r_B-r_A\approx(\mu_A-\mu_B)+(\varepsilon_B-\varepsilon_A).$$

Substituting these limits into the terms of eq. (SI1.25), which we analyze separately, we have

$$\begin{aligned}
r_B^2-r_A^2 =&(r_B-r_A)(r_B+r_A)\\
&\approx[(\mu_A-\mu_B)+(\varepsilon_B-\varepsilon_A)]\,[(\varepsilon_B-\mu_B)+(\varepsilon_A-\mu_A)]\\
&\approx\mu_B^2-\mu_A^2-2(\mu_B\varepsilon_B-\mu_A\varepsilon_A)+\varepsilon_B^2-\varepsilon_A^2\\
\delta\,[\,r_B-r_A+\mu_B-\mu_A\,]&\approx\delta(\varepsilon_B-\varepsilon_A),\\
r_B\mu_B-r_A\mu_A&\approx-\mu_B^2+\mu_A^2+(\mu_B\varepsilon_B-\mu_A\varepsilon_A)
\end{aligned}$$

and

$$a_m(r_A+\mu_A)(r_B+\mu_B)(r_B-r_A)\approx a_m\,\varepsilon_A\varepsilon_B(\varepsilon_B-\varepsilon_A+\mu_A-\mu_B).$$

Collecting,

$$\begin{aligned}
N_\mu =&\mu_B^2-\mu_A^2-2(\mu_B\varepsilon_B-\mu_A\varepsilon_A)+\varepsilon_B^2-\varepsilon_A^2+\delta(\varepsilon_B-\varepsilon_A)-\mu_B^2+\mu_A^2+(\mu_B\varepsilon_B-\mu_A\varepsilon_A)\\
&+a_m\,\varepsilon_A\varepsilon_B(\varepsilon_B-\varepsilon_A+\mu_A-\mu_B)\\
=&-(\mu_B\varepsilon_B-\mu_A\varepsilon_A)+\varepsilon_B^2-\varepsilon_A^2+\delta\varepsilon_B-\delta\varepsilon_A+a_m\,\varepsilon_A\varepsilon_B(\varepsilon_B-\varepsilon_A+\mu_A-\mu_B)\\
=&\varepsilon_B(\delta-\mu_B)-\varepsilon_A(\delta-\mu_A)+\underbrace{\varepsilon_B^2-\varepsilon_A^2}_{O(\varepsilon^2)}+\underbrace{a_m\,\varepsilon_A\varepsilon_B(\mu_A-\mu_B)}_{O(a_m\varepsilon^2)}+\underbrace{a_m\,\varepsilon_A\varepsilon_B(\varepsilon_B-\varepsilon_A)}_{O(a_m\varepsilon^3)}
\end{aligned}$$

Since $\mu_A<\mu_B$, in the limit $a_m\to\infty$, $\varepsilon_A$ is exponentially smaller than $\varepsilon_B$ because

$$\frac{\varepsilon_A}{\varepsilon_B}=e^{a_m(\mu_A-\mu_B)}\to 0\qquad\text{as } a_m\to\infty.$$

Therefore, we have

$$0 < \varepsilon_A \ll \varepsilon_B \ll 1.$$

Consequently,

$$\lim_{a_m \to 0} N_\mu \approx \varepsilon_B(\delta - \mu_B) - \varepsilon_A(\delta - \mu_A) + O(\varepsilon^2) + O(a_m\varepsilon^2) + O(a_m\varepsilon^3) > 0$$

because $\delta - \mu_B > 0$ and $\varepsilon_B \gg \varepsilon_A$. Thus, $\lim_{a_m \to \infty} \frac{\partial c_\mu}{\partial a_m} = 0$, yet for large but finite $a_m$, $\frac{\partial c_\mu}{\partial a_m} > 0$ when $\mu_i < \delta$.

When selection acts on post-maturity mortality, two regimes are possible:

1. when immature individuals experience a higher mortality rate than mature ones, $\mu < \delta$, the most common case in nature, the selection coefficient initially decreases with age at maturity until the threshold $a_m = 1/(\delta - \mu)$, and then increases. Therefore, the smaller the difference between immature and mature mortality rates, the larger the range over which the selection coefficient decreases with age at maturity.

2. when mature individuals experience a higher mortality rate than immature ones, $\mu > \delta$, the selection coefficient always decreases monotonically with age at maturity.

#### Supplementary Information 2: Description of the Individual-Based Model

We simulate adaptive evolution in a population facing a gradually changing environment using an individual-based model (IBM). The simulation proceeds in discrete time steps interpreted as units of absolute time. Three different scenarios are simulated: in the first, adaptation occurs via selection on fecundity, in the second via selection on pre-maturity survival, and in the third via selection on post-maturity survival.

To simulate a gradually deteriorating environment, we consider the optimal phenotype $\theta$ to shift at rate $\varepsilon$. Therefore, the change in the optimal phenotype in a time step $\Delta t$ equals

$$\Delta\theta = \varepsilon\,\Delta t.$$

In the model, each individual is characterized by their age $a$ and a quantitative trait $z$, which remains constant through life. Four individual processes are considered: aging, maturation, birth and death. At every time step, all individuals age by one unit. Individuals mature once their age reaches the age at maturity, $a_m$.

*Reproduction*

We consider clonal reproduction. Only mature individuals can reproduce. In the first scenario, fecundity is trait-dependent, following:

$$b(z) = \max\left(0, \beta e^{-\frac{(\theta - z)^2}{2\tau^2}} - b_s N\right). \tag{1}$$

This implies that an individual's birth rate $b$ equals $\beta$, the maximum per capita birth rate, when its trait $z$ is equal to the optimum phenotype $\theta$, and decreases as the difference between $z$ and $\theta$ increases. How fast the birth rate decreases with an increasing mismatch between the trait value and the optimum phenotype is determined by $\tau$. To illustrate this in a thermal gradient, the furthest the environmental temperature is from the thermal optimum of an individual, the lowest is its birth rate. In addition, the decrease in birth rate depends on the degree of specialization of the individual over the thermal gradient: a thermal specialist individual (small $\tau$) will have a larger reduction in birth rate with increasing mismatch between its trait value and the optimal phenotype than a thermal tolerant individual (large $\tau$). Similar trait-based approaches have been used by (*1*, *2*). Additionally, the birth rate decreases with increasing population size $N$ in a linear manner, where $b_s$ is the slope.

In the second and third scenario, an individual's birth rate is trait-independent, according to $b = \beta - b_s N$.

The birth rate $b$ determines whether an individual reproduces, and if so, how many offspring it produces in each time step. When $b \leq 1$, an individual produces one offspring with probability $b$ in a time step $\Delta t$. When $b > 1$, an individual produces an integer number of offspring equal to the floor of $b$, plus one additional offspring with probability equal to the fractional part of $b$. For example, if $b = 2.2$, this individual produces 2 newborns with probability 0.8 and 3 with probability 0.2. Newborns inherit the mother's trait value and have an age $a = 0$. Although mutation is implemented in the code, we disable this process by assuming a mutation rate of zero. This is because we focus on evolutionary responses from standing genetic variation only.

*Mortality*

The mortality rate differs among immature and mature individuals. In the second scenario, pre-maturity mortality is trait-dependent:

$$d_p(z) = \delta_{\min} + (\delta_{\max} - \delta_{\min})\left(1 - e^{-\frac{(\theta - z)^2}{2\tau^2}}\right) + d_s N. \tag{2}$$

Therefore, the pre-maturity mortality rate $d_p$ increases as the difference between the trait $z$ and the optimum phenotype $\theta$ increases, bounded between $\delta_{\min}$ and $\delta_{\max}$, and modulated by $\tau$. In addition, the mortality rate increases with increasing population size $N$ in a linear manner, where $d_s$ is the slope. In the two other scenarios, pre-maturity mortality is trait-independent, such that $d_p = \delta_{\min} + d_s N$, where $\delta_{\min}$ is a constant background mortality rate.

Similarly, in the third scenario, post-maturity mortality is trait-dependent, following

$$d_m(z) = \omega_{\min} + (\omega_{\max} - \omega_{\min})\left(1 - e^{-\frac{(\theta - z)^2}{2\tau^2}}\right) + d_s N, \quad (3)$$

where the parameters $\omega_{\min}$, $\omega_{\max}$, $\tau$ and $d_s$ denote analogous quantities to $\delta_{\min}$, $\delta_{\max}$, $\tau$ and $d_s$ in eq. 2. Conversely, in the first and second scenarios, post-maturity mortality is trait-independent, such that $d_m = \omega_{\min} + d_s N$, where $\omega_{\min}$ is a constant background mortality rate.

To determine which individuals die at each time step, we draw for each individual a random number from a uniform distribution on the interval [0,1]. If this number is smaller than the mortality rate $d_p$ or $d_m$, depending on the individual's life stage, this individual dies and is removed from the population.

We summarize the fecundity and mortality functions for each scenario in Table SI2.1.

Table SI2.1. Functions defining the demographic process in each scenario

| Demographic rate | Scenario 1 | Scenario 2 | Scenario 3 |
|---|---|---|---|
| Fecundity | $b(z) = \max\left(0, \beta e^{-\frac{(\theta-z)^2}{2\tau^2}} - b_s N\right)$ | $b = \beta - b_s N$ | $b = \beta - b_s N$ |
| Pre-maturity mortality | $d_p = \delta_p + d_s N$ | $d_p(z) = \delta_{\min} + (\delta_{\max} - \delta_{\min})\left(1 - e^{-\frac{(\theta-z)^2}{2\tau^2}}\right) + d_s N$ | $d_p = \delta_p + d_s N$ |
| Post-maturity mortality | $d_m = \delta_m + d_s N$ | $d_m = \delta_m + d_s N$ | $d_m(z) = \delta_{\min} + (\delta_{\max} - \delta_{\min})\left(1 - e^{-\frac{(\theta-z)^2}{2\tau^2}}\right) + d_s N$ |

*Model parameterization*

Because the evolutionary response can be suppressed when density regulation occurs via the same demographic rate that selection targets (*3*), the models alternatively implement density regulation via mortality in the fecundity selection scenario and via fecundity in the two survival selection scenarios. In other words, in the first scenario, we set $b_s$ to zero, whereas in the other two scenarios, we set $d_s$ to zero.

For trait-dependent mortality, the upper boundary of mortality $\delta_{\max}$ and $\omega_{\max}$ is set to be one order of magnitude higher than the lower boundary $\delta_{\min}$ and $\omega_{\min}$ (i.e. $\delta_{\max} = 10\ \delta_{\min}$ and $\omega_{\max} = 10\ \omega_{\min}$), ensuring a substantial but biologically realistic range of mortality variation.

For simulations of the simplified demographic framework, in which only age at maturity varies (figure 3), we use the parameters shown in Table SI2.2. The values of the demographic parameters are within the range of variation in natural populations, as reported in the dataset compiled by Healy et al. (*4*).

Table SI2.2. Model parameters used in figure 3

| **Parameters** | **Symbol** | **Value** | | |
|---|---|---|---|---|
| | | Scenario 1 | Scenario 2 | Scenario 3 |
| Environmental parameters | | | | |
| Rate of degradation | $\varepsilon$ | 0.0001 | 0.00005 | 0.0002 |
| Demographic parameters | | | | |
| Maximum birth rate | $\beta$ | 0.05 | 1 | 0.8 |
| Baseline pre-maturity mortality | $\delta_{\min}$ | 0.005 | 0.005 | 0.005 |
| Baseline post-maturity mortality | $\omega_{\min}$ | 0.0005 | 0.0005 | 0.0005 |
| Coefficient for the effect of density-dependence on fecundity | $b_s$ | 0 | 0.001 | 0.001 |
| Coefficient for the effect of density-dependence on mortality | $d_s$ | 0.00001 | 0 | 0 |

For simulations informed by empirical demographic rates (figure 4), the optimal-condition parameters $\beta$ (maximum fecundity), $\omega_{\min}$ and $\delta_{\min}$ (baseline mortalities) are estimated from the dataset compiled by Healy et al. (*4*). This dataset includes demographic information for 279 wild animal populations representing 120 species, drawn from the COMADRE database (version 2.0.1.1). These populations were specifically selected because they correspond to unmanipulated wild populations and therefore reflect natural demographic processes. Two populations (*Sterna hirundo* and *Ovis canadensis*) exhibit zero pre-maturity mortality, an unrealistic mortality rate. These were therefore excluded, resulting in a final dataset of 277 populations used for parameterization, whose demographic rates are shown in figure S2. The density-dependent birth and death components $b_s$ and $d_s$ are taken as in the simplified demographic framework.

Table SI2.3. Model parameters used in figure 4

| Parameters | Symbol | Value<br>Scenario 1 | Scenario 2 | Scenario 3 |
|---|---|---|---|---|
| Environmental parameters | | | | |
| Rate of degradation | $\varepsilon$ | 0.0005 | 0.00002 | 0.0001 |
| Demographic parameters | | | | |
| Maximum birth rate | $\beta$ | Species-specific demographic information from Healy et al. (see figure S2) | | |
| Baseline pre-maturity mortality | $\delta_{\min}$ | Species-specific demographic information from Healy et al. (see figure S2) | | |
| Baseline post-maturity mortality | $\omega_{\min}$ | Species-specific demographic information from Healy et al. (see figure S2) | | |
| Coefficient for the effect of density-dependence on fecundity | $b_s$ | 0 | 0.001 | 0.001 |
| Coefficient for the effect of density-dependence on mortality | $d_s$ | 0.00001 | 0 | 0 |

*Model initialization*

Each simulation starts in an environment where the population is well adapted, meaning that its mean trait equals the optimal phenotype. The starting population is initialized at its stable age distribution, and individual trait values are drawn from a normal distribution with mean equal to the optimum and a standard deviation $\sigma$ of 0.1. We use this standard deviation to represent the amount of standing genetic variation available for evolution. Because traits in our model are clonal and lack environmental variation, phenotypic variance corresponds directly to additive genetic variance. A standard deviation of 0.1 therefore implies an additive genetic variance of $0.1^2$=0.01, which lies within the range of empirical estimates of additive genetic variance in fitness components (0.003–0.5) reported for 19 natural populations (*5*).

*Quantifying fitness*

At the end of each simulation, we compute the initial, final fitness without evolution, and final fitness with evolution. As a measure of fitness, we use the intrinsic growth rate $r$, defined implicitly by the Euler-Lotka equation (eq. 1). Solving the equation yields:

$$r = \frac{1}{a_m} W\left(a_m b\, e^{a_m(d_m - d_p)}\right) - d_m,$$

where $W(\cdot)$ denotes the Lambert W function. To isolate the contribution of an individual's vital rates to fitness, we evaluate $r$ under density-independent conditions, setting the density-dependent birth and death components $b_s$ and $d_s$ to zero.
